# BRI1-mediated removal of seed coat H3K27me3 marks is a brassinosteroid-independent process

**DOI:** 10.1101/2023.12.07.569203

**Authors:** Rishabh Pankaj, Rita B. Lima, Guan-Yu Luo, Sinah Ehlert, Gerardo del Toro-de León, Heinrich Bente, Pascal Finger, Hikaru Sato, Duarte D. Figueiredo

## Abstract

Seed development in angiosperms starts with double fertilization, where two paternal sperm cells fertilize the maternal gametes. This leads to the formation of the embryo and of the endosperm. These fertilization products are enveloped by the maternally-derived seed coat, the development of which is inhibited prior to fertilization by the Polycomb Repressive Complex 2 (PRC2). This complex deposits the repressive histone mark H3K27me3, whose removal is necessary for seed coat formation. Here, we show that JUMONJI-type (JMJ) histone demethylases are expressed in the seed coats of *Arabidopsis thaliana* (Arabidopsis) and are necessary for its formation. We propose that JMJ activity is coupled to Brassinosteroid (BR) function, as BR effectors physically recruit JMJ proteins to target loci. Consistent with this, we show that loss of BR biosynthesis and signaling leads to seed coat defects, and that loss of the main BR receptor, BRI1, results in H3K27me3 hypermethylation. Moreover, our data points to BRI1 mediating H3K27me3 removal independently of BRs, while a different receptor, BRL3, likely regulates seed coat formation in a BR-dependent manner. We thus propose a model where seed coat development relies on canonical and non-canonical functions of BR receptors.

## Introduction

Changes in chromatin accessibility driven by epigenetic modifications, including DNA methylation and histone marks, modulate gene expression and thus shape plant and animal development. Plants have extensive systems in place to control the deposition and removal of these epigenetic marks on their chromatin (Hemenway and Gehring, 2023). Among such marks, H3K27me3 is a critical histone modification that induces gene repression and chromatin compaction (Mozgova et al., 2015). This mark is deposited by Polycomb Group proteins (PcG), which form multimeric complexes known as POLYCOMB REPRESSIVE COMPLEXES (PRC). This includes complexes of the PRC1 and PRC2 types, the latter being responsible for H3K27me3 deposition. In the model system *Arabidopsis thaliana* (Arabidopsis), there are three PRC2 complexes, each with its unique collection of components and roles in plant growth: EMBRYONIC FLOWER (EMF), VERNALIZATION (VRN), and FERTILIZATION-INDEPENDENT SEED (FIS). The EMF- and VRN-PRC2s are specific to the sporophytic generation, while FIS-PRC2 is specific to the gametophyte and to one of its products, the endosperm (Mozgova et al., 2015).

The developing seed of an angiosperm contains three genetically distinct structures: embryo, endosperm, and seed coat. While the formation of the first two is directly linked to the fertilization of the egg cell and of the central cell, the seed coat, which derives from the ovule integuments, is not a direct product of fertilization. This has two implications: 1) seed coat development is blocked prior to fertilization, and 2) the ovule integuments require a signal originating from the fertilization products, to drive seed coat formation. Indeed, the development of the seed coat is actively blocked prior to fertilization by the sporophytic PRC2s, EMF and VRN (Roszak and Köhler, 2011). Moreover, the hormone auxin is the signal coupling fertilization to seed coat development (Figueiredo et al., 2016). Following fertilization, the endosperm produces auxin (Figueiredo et al., 2015), which is then transported to the integuments, where it removes the PRC2s, allowing for seed coat development (Figueiredo et al., 2016). In summary, sporophytic PRC2s are responsible for restricting seed coat formation before fertilization and this block is lifted by auxin. This is supported by the observation that mutants lacking sporophytic PRC2 activity, and thus have reduced levels of H3K27me3, produce fertilization-independent (or autonomous) seed coats (Figueiredo et al., 2016; Roszak and Köhler, 2011). This is the case for mutants lacking the PRC2 components SWINGER (SWN), CURLY LEAF (CLF), VERNALIZATION 2 (VRN2) and EMBRYONIC FLOWER 2 (EMF2). Genes encoding these PRC2 subunits are downregulated upon fertilization or upon exogenous application of auxin (Figueiredo et al., 2016). Concurrently, treating ovules with exogenous auxin leads to autonomous seed coat formation, as auxin removes PRC2 function (Figueiredo et al., 2016). However, the auxin-drived removal of PRC2 does not explain how seed coat genes become active, given that the H3K27me3 marks should be stable. Importantly, seed coat growth is not driven by cell division but only by cell elongation (Figueiredo et al., 2016), meaning that dilution of the marks is unlikely. Therefore, the H3K27me3 marks must likely be actively removed following fertilization, but the enzymes responsible for this process are still unknown.

Histone demethylases, such as those containing JumonjiC domains (JmjC), can remove histone marks such as H3K27me3. Arabidopsis has 21 predicted JmjC proteins that can be categorized into five classes based on the architecture of their protein domains (Crevillén, 2020). While not all members have been fully studied, they include potential H3K9me2, H3K36me3, H3K4me3, and H3K27me3 demethylases. In Arabidopsis, five H3K27me3 demethylases have been identified so far, including two JmjC domain-only proteins, JUMONJI 30 (JMJ30/AtJMJD5) and JUMONJI 32 (JMJ32), as well as the C2H2-type zinc-finger (ZnFn)-containing JmjC proteins, EARLY FLOWERING 6 (ELF6/JMJ11) and RELATIVE OF ELF6 (REF6/JMJ12), and their closely related JUMONJI 13 (JMJ13) (Crevillén et al., 2014; Cui et al., 2016; Gan et al., 2014; F. Lu et al., 2011; S. X. Lu et al., 2011; Yan et al., 2018). However, it is possible that additional H3K27me3 demethylases are yet to be discovered. The three major H3K27me3 demethylases REF6, ELF6 and JMJ13 have been shown to be important for reproductive processes. REF6 was shown to be necessary for suppression of the seed dormancy (Chen et al., 2020), as well as for seed germination (Pan et al., 2023; Sato et al., 2021; Wang et al., 2023). ELF6 and JMJ13, on the other hand, have been shown to control carpel growth in an antagonistic matter (Keyzor et al., 2021). ELF6 was also shown to be expressed in the mature ovules and in the embryo (Crevillén et al., 2014; Yang et al., 2016). Finally, JMJ13 has been shown to be necessary for genome-wide H3K27me3 demethylation in the pollen (Borg et al., 2020).

Although JMJs play key roles in epigenetic reprogramming, these proteins often must be recruited to their target loci by transcription factors (TF). This includes TFs involved in brassinosteroid (BR) signaling, like BRASSINAZOLE-RESISTANT 1 (BZR1) and BRI1-EMS-SUPPRESSOR 1 (BES1) (Li et al., 2018; Yu et al., 2008). As hormones of steroid nature, BRs serve a variety of roles in lant development (Manghwar et al., 2022). They control cell division, stem cell maintenance, vascular development, cell elongation, root growth and floral transition, among other processes (Fàbregas and Caño-Delgado, 2014; Lv et al., 2018; Singh and Savaldi-Goldstein, 2015; Vukašinović et al., n.d.; Wang et al., 2020). Moreover, BRs have been extensively implicated in reproductive processes, including in seed development (Lima and Figueiredo, 2024). And this is the case in several species. For example, rice BR biosynthesis and signaling mutants produce smaller seeds (Hong et al., 2005; Morinaka et al., 2006; Tanabe et al., 2005). The same is true for *Vicia faba* (Fukuta et al., 2006) and pea (Nomura et al., 2007). In Arabidopsis, the BR-deficient mutant *dwf5* also produces small seeds (Choe et al., 2000), and reduction of endogenous BR levels by ectopic expression of the P450 monooxygenase family gene *CYP72C1* also results in general dwarfed organs and small seeds (Takahashi et al., 2005). In addition to this, the BR-deficient mutant *deetiolated 2* (*det2*) and the BR-insensitive mutant *bri1-5* (a weak allele of the main BR receptor BRASSINOSTEROID INSENSITIVE 1) also make seeds that are smaller and shaped differently than the respective wild-type (WT) (Jiang et al., 2013). These seed phenotypes of *det2* can be partly rescued by exogenous BR, supporting the positive regulatory role for BRs in seed growth (Jiang et al., 2013). This effect of BRs on the development of Arabidopsis seeds was proposed to be determined by the direct regulation of genes involved in the HAIKU (IKU) pathway, which determines endosperm size, by the BR effector BRASSINAZOLE INSENSITIVE 1 (BZR1) (Garcia et al., 2003; Luo et al., 2005; Wang et al., 2010; Zhou et al., 2009).

Although BRs have been implicated in regulating seed growth, the underlying molecular mechanisms are still poorly understood. Importantly, as alluded to above, the BR effector BES1 was shown to regulate the expression of target genes by recruiting the JMJ histone demethylases ELF6 and REF6 (Yu et al., 2008). Interestingly, BES1 interacts physically with both ELF6 and REF6 (Yu et al., 2008), while BZR1 interacts with ELF6 but not REF6 (Li et al., 2018). We thus hypothesized that JMJ and BR function could be necessary to remove H3K27me3 marks from the integuments, allowing the seed coat to develop after fertilization. Indeed, here we show that BR and JMJ mutants show seed coat defects. Consistent with our hypothesis that BRs are necessary for H3K27me3 removal, we show that some BR mutant phenotypes correlate with H3K27me3 hypermethylation and are rescued by loss of PRC2 function in the integuments. Moreover, we uncover a dual role for BR regulation of seed coat growth, mediated by the main BR receptor BRI1 and by one of its close homologues BRI-LIKE 3 (BRL3).

## Results

### REF6 and ELF6 H3K27me3 demethylases are expressed in the seed coat

Previous research indicates that seed coat development is blocked by H3K27me3 marks, which are deposited by sporophytic PRC2s (Figueiredo et al., 2016; Roszak and Köhler, 2011). Auxin-mediated removal of the PRC2s is therefore necessary for seed coat formation following fertilization (Figueiredo et al., 2016). However, PRC2 removal alone does not explain seed coat formation, because the H3K27me3 marks should be stable in the non-dividing seed coat cells. If this is true, then ovules of mutants partly lacking PRC2 function, and therefore depleted in H3K27me3, should be more responsive to exogenous auxin, and should develop larger autonomous seed coats, when compared to the WT. Indeed, we observed that the sporophytic PRC2 mutant *swn clf*/*+* produces larger autonomous seeds than Col-0 after treatments with 100 µM of the synthetic auxin 2,4-Dichlorophenoxyacetic acid (2,4-D; **Fig. S1**). This observation supports the hypothesis that removal of H3K27me3 marks in the seed coat after fertilization is an essential step for seed coat growth. We then hypothesized that H3K27me3 marks should be enzymatically removed from the integument cells following fertilization. This process can be carried out by JMJ-type histone demethylases. Therefore, we analyzed previously published seed-specific transcriptomic datasets to test if JMJ encoding genes are expressed in the seed coat (**Fig. S1**). Indeed, several genes encoding H3K27me3 demethylases are predicted to be expressed during seed coat development, including *ELF6*, *JMJ13* and *JMJ30* (Belmonte et al., 2013). Unfortunately, the microarray dataset that we used did not contain a probe for *REF6* .

Because in Arabidopsis the removal of H3K27me3 marks is carried out by three main H3K27me3 demethylases: REF6, ELF6 and JMJ13, we decided to focus our analyses on these members of the JMJ family. Through analysis of a *REF6::REF6:GFP* reporter in a *ref6c* mutant background (Yan et al., 2018), we found that REF6 is expressed in the integuments of unfertilized ovules, as well as in seed coats of 1 DAP seeds (**Fig. 1A**). At these stages of development, no REF6 expression was seen in the gametophyte or in the early endosperm or zygote. We confirmed these observations using a line expressing *REF6::GUS*, and consistently observed GUS activity in sporophytic and zygotic tissues of developing seeds as well as a strong expression in anthers (**Fig. S2**). Regarding ELF6, it was previously found to be expressed in mature ovules as well as in developing embryos (Crevillén et al., 2014). We analyzed a reporter line expressing *ELF6::GUS* and indeed observed GUS activity in the sporophytic tissues of developing seeds (**Fig. 1B**). Finally, we analyzed JMJ13 transcriptional (*JMJ13:GFP*) and translational (*JMJ13:JMJ13-GFP*) reporters and we did not observe any *JMJ13* expression in seeds (**Fig. S2**). To test if these reporter lines recapitulated the published expression patterns, we checked their expression in developing anthers, as JMJ13 has been shown to be expressed in pollen (Borg et al., 2020). Indeed, fluorescence was observed in mature pollen grains (**Fig. S2**), confirming that the reporters are functional and that JMJ13 is likely not expressed in seeds. These expression results indicate that among the three main H3K27me3 demethylases, REF6 and ELF6 are the ones with the strongest expression in seeds, whereas together with REF6, JMJ13 is mostly expressed in pollen.

**Figure 1.**
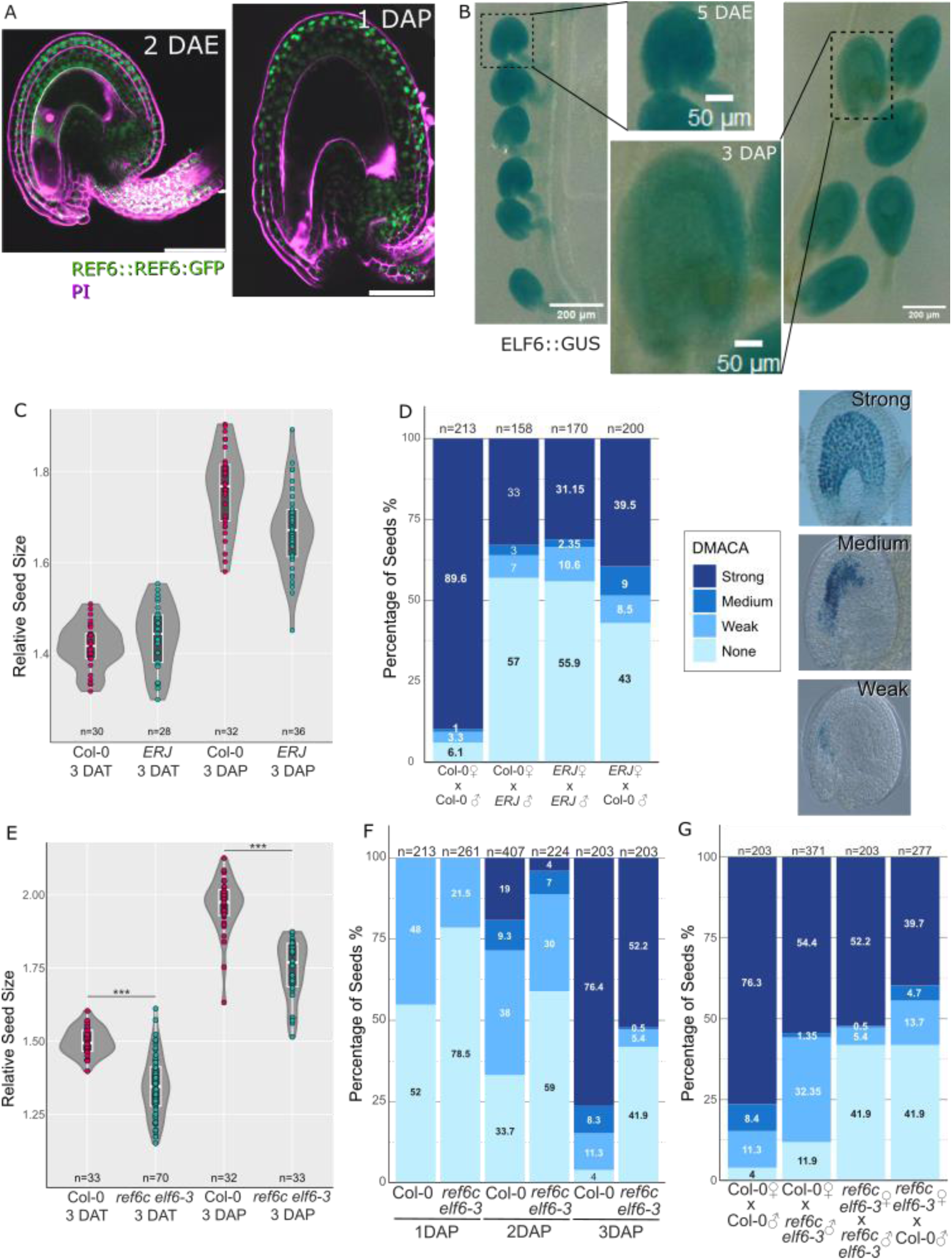
JMJ activity is necessary for seed coat formation. (**A**) Expression of *REF6::REF6-GFP* in *ref6c* in an unfertilized ovule at 2 days after emasculation (DAE; top) and 1 day after pollination (DAP; bottom). Scale bars indicate 50 µm. Magenta is propidium iodide. (**B**) Expression of *ELF6:GUS* in unfertilized ovules and developing seeds at 3 DAP. (**C**) Autonomous and sexual seed size of *elf6 ref6c jmj13 (ERJ*) and the respective WT. The relative seed size was calculated as a ratio between the perimeter of each seed and the average perimeter of unfertilized ovules of their respective genotype. (**D**) DMACA staining of *ERJ* x Col-0 reciprocal crosses at 3 DAP. The staining was classified in four categories: examples shown on the right hand-side for the three stained categories. (**E**) Autonomous and sexual seed size of *ref6c elf6-3* and the respective WT. The relative seed size was calculated as a ratio between the perimeter of each seed and the average perimeter of unfertilized ovules of their respective genotype. *** Differences are significant for 0.001>p (ANOVA). (**F-G**) DMACA staining of *ref6c elf6-3* at 1, 2 and 3 DAP (**E**), and of *ref6c elf6-3* x Col-0 reciprocal crosses (**F**).

### Mutants for JMJ type H3K27me3 demethylases have seed coat development defects

In order to test whether JMJ histone demethylases are responsible for H3K27me3 demethylation in the seed coat, we assessed seed coat formation in *ref6-1, elf6-3* and *jmj13* single mutants. We did this both using sexual and asexual (or autonomous) seeds, using seed size as a proxy for seed coat expansion. Shortly, we compared the size of sexual seeds 3 days after pollination (referred to as 3 DAP) and/or of autonomous seeds 3 days after 100 µM auxin (2,4-D) treatment (referred to as 3 DAT). We selected this time point, because in these early stages of seed development the expansion of the seed is purely driven by seed coat growth and its interactions with the endosperm. For instance, the size of a mature Arabidopsis seed is also determined by the embryo, and therefore we used a timepoint in which the embryo size did not impact on our measurements. The reasoning for analyzing autonomous seeds, as produced via exogenous auxin applications, in addition to sexual seeds, was to test if auxin-induced seed coat formation is specifically impaired in *jmj* mutants.

At 3 DAP we observed that seeds of the single r*ef6c, elf6* and *jmj13* mutants were of similar sizes to corresponding WT (**Fig. S3**). We also did not find consistent defects in the size of autonomous seeds at 3 DAT (**Fig. S3**). This indicates that the *jmj* single mutants do not show any major seed coat initiation defects. Since ELF6, REF6 and JMJ13 share significant homology, it is possible that they redundantly control seed coat formation. Consequently, we analyzed higher order mutants for these genes. We obtained and analyzed a published *elf6 ref6c jmj13 (ERJ)* triple mutant (Yan et al., 2018). Plants carrying these three mutant alleles are dwarf, exhibit delayed flowering time relative to the WT, and show bent siliques, as previously described (**Fig. S3 and S4**) (Yan et al., 2018). This was not observed in the single mutants, indeed supporting the idea of functional redundancy between these JMJs. We then conducted a similar comparative analysis of the sizes of sexual and autonomous seeds. Surprisingly, and contrary to our expectations, we found that both sexual and autonomous seeds of the triple mutant were slightly but significantly larger than those of the wild type (**Fig. S3**). However, we noted that the unfertilized/mock ovules of the triple mutant were also significantly bigger than those of the wild type (see mock-treated samples in **Fig. S3**). And thus, when normalizing for the size of unfertilized ovules, we detected a slight, but not statistically significant, reduction in relative seed coat growth in the *ERJ* mutant compared to the WT (**Fig. 1C**). Therefore, we used an alternative approach to assess seed coat formation: we performed DMACA (p-dimethylaminocinnamaldehyde) staining of seeds at 1, 2 and 3 DAP. This dye stains the proanthocyanidins (PAs) produced in the endothelium of the seed coat, and can therefore be used as a visual marker for seed coat development (Debeaujon et al., 2003). Supporting our hypothesis, we observed that seeds of the *ERJ* triple mutant showed a striking delay in the accumulation of proanthocyanidins, when compared to the WT (**Fig. 1D**). This was obvious in all three time points assessed, and fits with our expectation that JMJ function is necessary for seed coat development.

Our analyses also revealed that the siliques of *ERJ* contained many immature ovules or aborted seeds (**Fig. S3 and S4**). There could be several reasons for this: 1) a post-fertilization effect on seed viability; 2) compromised pollen function; or 3) compromised ovule viability. To test if any of these was true, we did DMACA staining of 3 DAP seeds of reciprocal crosses of *ERJ* × Col-0 (**Fig. 1D**). This analysis revealed several reproductive defects in the *ERJ* mutant, which results in a reduced number of fertilized seeds. Specifically, we observed that in *ERJ* × Col-0 (by convention the maternal parent is indicated first), 43% of the total seeds in the siliques did not stain with DMACA, even after 3 days. Upon closer inspection we observed that many of these ovules were not fully mature (**Fig. S3**), which led to a lower number of fertilized seeds. Additionally, in Col-0 × *ERJ* crosses we again observed that many seeds did not stain with DMACA, and approximately 57% of ovules were in fact unfertilized. This is likely due to defects in *ERJ* pollen viability. Nevertheless, in *ERJ* × Col-0 crosses, where the seed coat is derived from the triple mutant, a significant proportion of viable seeds (∼17.5%) were delayed in seed coat development, as indicated by less DMACA staining compared to their wild-type counterparts (WT; categories “Medium” and “Weak” in **Fig. 1D and S3**). This fits our hypothesis that these JMJs are necessary for seed coat development.

However, despite these findings, the pleiotropic defects of the triple mutant complicated its analysis. To counter this, we analyzed less strong double mutants, and phenotyped seeds of *ref6-1 elf6-3*, *elf6-3 jmj13* and *jmj13 ref6-1*. None of the three double mutants showed any visible vegetative growth phenotypes. Additionally, when assessing seed size at 3 DAP in the three double mutants, we observed that all three produced seeds that were slightly smaller than the WT ones (**Fig. S3**). Because JMJ13 does not seem to be expressed in seeds, as we describe above, we further focused on ELF6 and REF6 as potential regulators of seed coat development. In the double mutant analysis of **Fig. S3** we used *ref6-1*, which is a knock-down of REF6.

Therefore, we obtained and analyzed the stronger *ref6c elf6* mutant, where *ref6c* is a CRISPR-knock-out allele of *REF6* (Yan et al., 2018). Unlike *ref6-1 elf6-3*, the stronger *ref6c elf6* mutant had visible vegetative growth defects: the plants displayed dwarfed growth patterns, similar to those seen in the *elf6 ref6c jmj13* mutants, but did not exhibit their characteristic bent silique phenotype (**Fig. S3 and S4**). Importantly, the *ref6c elf6* double mutant produced young seeds that were significantly smaller than the WT ones, as is expected for mutants with seed coat initiation defects (**Fig. 1E**). This was true both for sexual seeds and for those obtained via exogenous auxin applications. Moreover, the *ref6c elf6* double mutant seeds showed seed coat defects when stained with DMACA (**Fig 1F-G**): *ref6c elf6* produced a significant number of seeds that did not stain with DMACA and, unlike in the WT, the staining did not increase as much in the mutant seeds as they developed (**Fig. 1F**). In fact, around 42% of the *ref6c elf6* seeds did not stain with DMACA even after 3 DAP. To further confirm the origin and nature of the defect, we performed reciprocal crosses of *ref6c elf6* with the WT (**Fig. 1G**). Interestingly, in the case of Col-0 X *ref6c elf6,* the pollen defects seem to be reduced when compared to the *ERJ* triple mutant (**Fig. S3**). This is in line with JMJ13 being specifically expressed in pollen (**Fig. S2**). In addition to these findings, it is important to note that *ref6c elf6* mutants still exhibited a decrease in the overall seed set, with only around 30 viable seeds per silique, when compared to around 50 seeds produced by the WT (**Fig. S3**). A reduced seed set normally correlates with larger individual seed size. Therefore, although we observed seed coat development defects in *jmj* mutants, it is possible that the full extent of the seed coat defects is partially masked by the low seed set of these lines.

In conclusion, JMJ function is required for seed coat formation. The JMJ H3K27me3 demethylases ELF6 and REF6 are expressed in the seed coat, and mutations in the respective genes lead to delayed seed coat growth and in accumulation of PAs. Additionally, loss of JMJ function also compromises ovule and pollen development, leading to reduced seed sets.

### JMJ function represses seed growth at later stages of development in a zygotic manner

Although our data supports a role for H3K27me3 demethylases in promoting seed coat initiation, we were surprised to observe that the *jmj* mutant seeds were actually larger at maturity compared to the respective WT. We tested this in several mutant alleles of *ref6* and *elf6*, as well as for the higher order mutants *elf6-3 ref6c* and *ERJ* (**Fig. 2A and Fig. S4**). Although in the case of the higher order mutants this increase in individual seed size could be due to the reduced seed set, as mentioned above, *ref6* single mutants also produced larger seeds at maturity (**Fig. 2A-B and S4**), although their seed set was comparable to that of the WT (**Fig. S3**). This suggests that JMJ function promotes early seed growth, but represses it at later stages of development. Because REF6 has been shown to have functions in endosperm development (Sato et al., 2021), we hypothesized that the increased size of *ref6* mature seeds could be due to a zygotic effect. To test this, we complemented the *ref6* mutant with constructs driving *REF6* expression under 1) its native promoter, 2) an embryo-specific promoter (*TPS1*) and 3) an endosperm-specific promoter (*EPR1*), as previously described (Sato et al., 2021). As expected, the mature seed size was restored to WT levels when *ref6* was complemented with *REF6::REF6:GFP* (**Fig. 2C**). Importantly, we also observed a partial but significant rescue of the *ref6* mature seed size phenotype in lines expressing *TPS1::REF6* and *EPR1::REF6* (**Fig. 2D**). This suggests that the increased size of *jmj* mutant seeds is, at least in part, due to zygotic effects of the embryo and the endosperm. If this is true, then REF6 should target genes that are specifically expressed in all three seed tissues. We thus searched for REF6 binding sites in embryo-, endosperm- and seed coat-specific genes, based on published datasets (Belmonte et al., 2013; Cui et al., 2016). Indeed, 326 genes bearing at least four REF6 binding motifs CTCTGYTY in their vicinity are specifically expressed in the three seed tissues (**Fig. 2E** and **S5**). Interestingly, there is little overlap between REF6 targets in those tissues, suggesting that REF6 controls different biological processes in the different tissues, and at different developmental timepoints. Thus, our data points to a sporophytic function of JMJs at early stages of seed development, promoting seed coat formation, and to a zygotic function at later stages of seed development, restricting embryo and endosperm growth. Consistent with this, *REF6* is strongly expressed in the seed coat in the first days of seed development, but its expression decreases in this tissue as the seeds develop (**Fig. S2**). By day 6 after pollination (6 DAP in **Fig. S2**) *REF6:GUS* expression is mostly absent from the seed coat and mostly detectable in the zygotic products.

**Figure 2.**
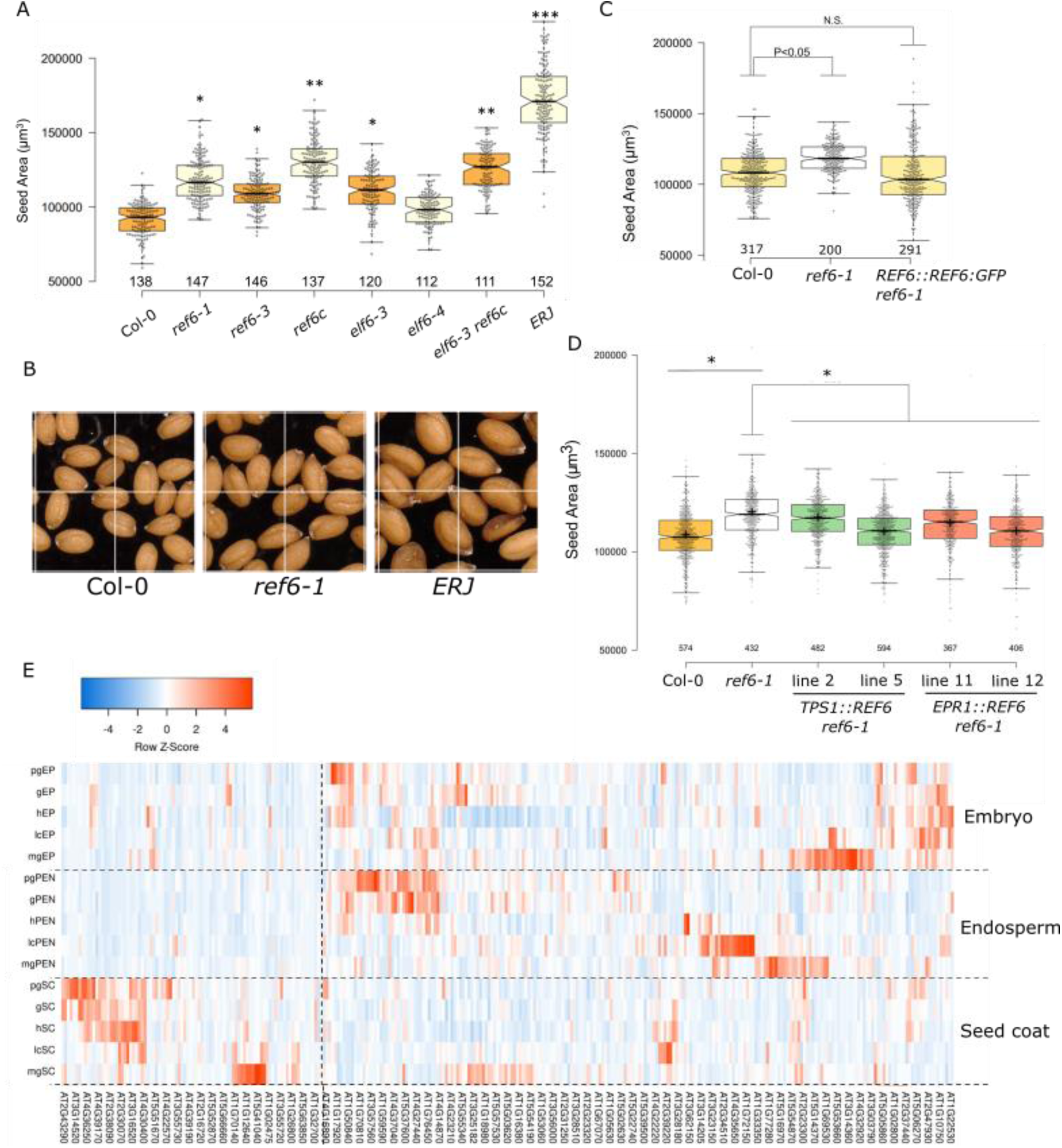
Zygotic effect of JMJs in mature seed growth. (**A**) Seed area of mature seeds of WT, *ref6* and *elf6* single mutants, and higher order JMJ mutants. Examples of seeds can be seen in (**B**) and in **Fig. S4**. (**C-D**) Mature seed area of WT, *ref6-1* and *ref6-1* complemented with the endogenous *REF6::REF6:GFP* (**C**), or with embryo (*TPS1*) or endosperm (*EPR1*) specific promoters (**D**). Two independent transgenic lines are shown. * indicates statistical significance for p<0.01 (Tukey multiple comparison) (**E**) Relative expression of genes carrying predicted REF6 binding sites, specifically expressed in the embryo proper (upper panel), peripheral endosperm (middle panel) and seed coat (lower panel). Extended dataset can be found in **Fig. S5**. The seed stages indicated are: pg, pre-globular; g, globular; h, heart; lc, linear cotyledon; and mg, mature green.

### BR mutants are defective in seed coat development

Given our observation that ELF6 and REF6 are redundantly required for seed coat formation, and given that both H3K27me3 demethylases have been shown to interact with the BR effectors BES1 and BZR1, we hypothesized that BR function could be required in the seed coat for H3K27me3 removal. We checked previously published datasets (Belmonte et al., 2013), and indeed genes involved in BR biosynthesis and signaling are predicted to be expressed in the seed coat (**Fig. 3A**). We recently confirmed that this is the case and all known enzymes involved in BR biosynthesis, as well as the BR receptor BRI1, co-receptor BAK1 and effectors BZR1 and BES1 are all specifically expressed in the early seed coat (Lima et al., 2023).

**Fig. 3.**
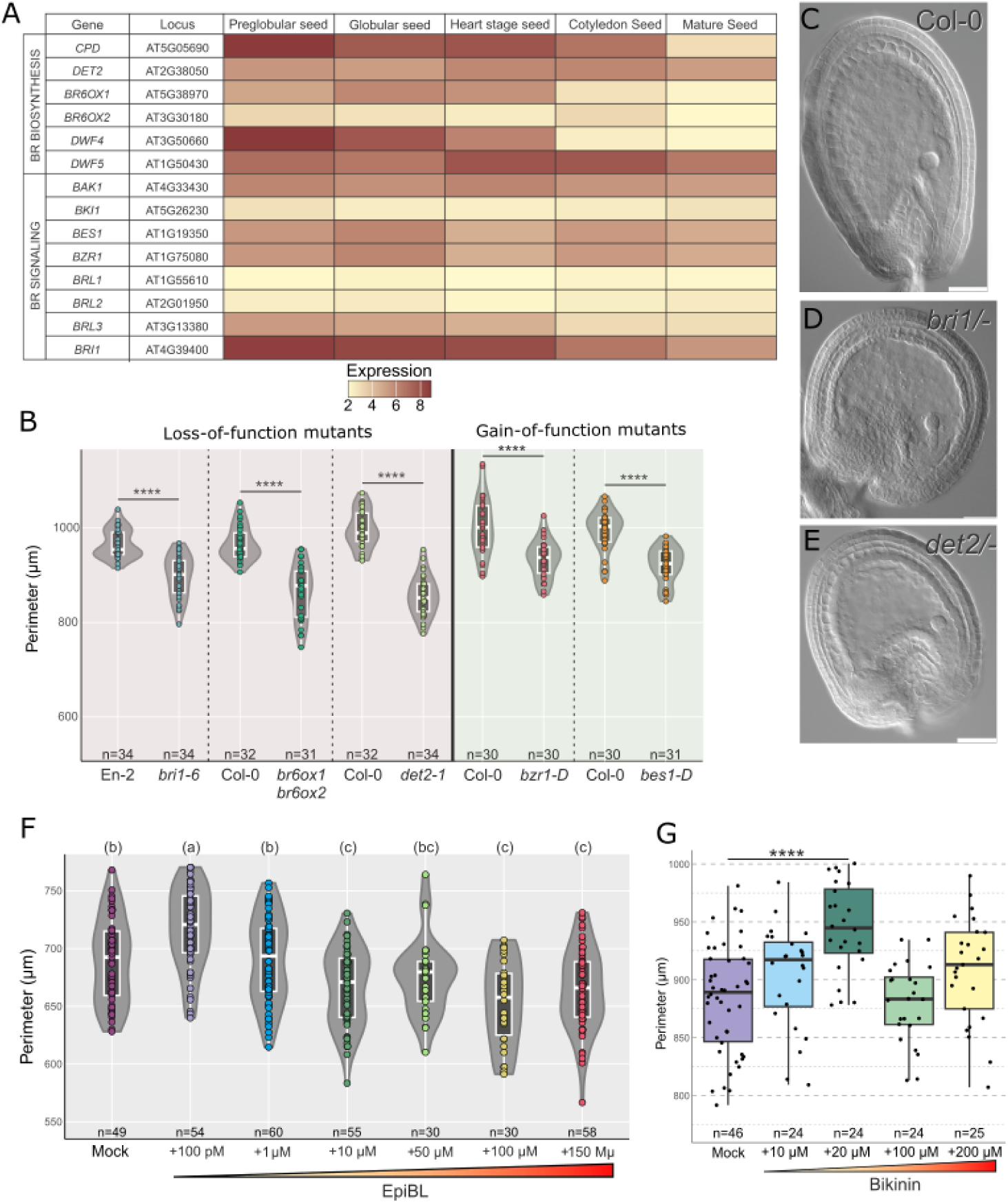
BR levels impact on seed coat development. (**A**) Relative expression of genes involved in BR biosynthesis and signaling in the seed coat, at different stages of seed development (indicated above), as determined in (Belmonte et al., 2013). (**B**) Perimeter of seeds at three DAP for loss- and gain-of-function BR mutants. The morphology of the seeds can be seen in **C-E**, for WT, *bri1* and *det2*. Scale bars indicate 50 μM. (**F-G**) Perimeter of 3 DAT autonomous seeds after exogenous application of 100 μM 2,4-D and varying concentrations of epi-brassinolide (EpiBL; **F**) or bikinin (**G**). **** represents p-value <0.0001 (Anova, **B**, **G**). For **F**, the letters on top indicate statistical significance for p-value <0.01 (Anova).

If our hypothesis is true, that BR signaling is required for H3K27me3 removal via JMJ function during seed coat formation, then BR-related mutants should show defects in seed coat development. Thus, we examined previously published BR mutants for seed defects. We tested loss-of-function mutants for components of BR signaling and biosynthesis. Indeed, we observed that mutants lacking the main BR receptor BRASSINOSTEROID INSENSITIVE 1 (BRI1) form seeds that are significantly smaller than those of WT at 3 DAP (**Fig. 3B-D**). The same is true for mutants impaired in BR biosynthesis like *br6ox1 br6ox2* and *det2-1* (*brassinosteroid-6-oxidase* and *deetiolated 2*; **Fig. 3D,E**). Similar to what we did for the JMJ mutants, we also analyzed the size of auxin-induced autonomous seeds. Again, we observed that all tested BR mutants initiate smaller autonomous seed coats, when compared to the WT (**Fig. S6**). For further experiments we selected one BR biosynthetic mutant, *det2-1*, and one signaling mutant, *bri1-6*. Importantly, the size of unfertilized ovules of *det2* and *bri1* plants was not significantly different from that of the wild type (**Fig. S6**), which signifies that the delay in seed growth is due to pathways that are activated after fertilization and not to initial ovule size. Overall, since the expression of BR genes is restricted to the seed coat, the smaller seed size in these mutants is likely a result of defects in seed coat development and not from a non-cell autonomous effect of the endosperm or embryo. Further evidence of this was obtained by expressing PHYB ACTIVATION TAGGED SUPPRESSOR 1 (BAS1), a protein responsible for degrading bioactive BRs (Turk et al., 2005), specifically in the seed coat. We observed that 4 out of 9 independent transgenic lines expressing the construct *KLUH:BAS1* produced seeds smaller than the WT control (**Fig. S6**). The promoter of *KLUH* is specific to the sporophytic tissues of the seed (Adamski et al., 2009). The effect was not as strong as what we observed for BR mutants, which is likely the result of the *KLUH* promoter not being expressed in all integument layers.

To further validate these results, we treated WT seeds with propiconazole, a known BR biosynthesis inhibitor (Hartwig et al., 2012). We applied 200 µM of propiconazole 6 h after pollination, to allow fertilization to take place. We observed that 3 DAP seeds treated with propiconazole were smaller than mock treated seeds (**Fig. S6**). This further confirms that BR function is necessary for seed coat growth.

### Exogenous BRs show a dose-dependent effect on seed size

We then tested the effects of constitutive BR biosynthesis or signaling in seed coat growth. For this, we analyzed several gain-of-function BR mutants. Our expectation was that those mutants would produce larger seed coats, when compared to the WT. We tested the mutant *dwf4-5D*, in which a T-DNA carrying a *CaMV35S* enhancer is inserted in the promoter of the BR biosynthesis gene *DWARF4* (*DWF4*), resulting in a constitutive production of BRs (Kim et al., 2013). Our results showed that the 3 DAP seeds of this mutant were significantly larger than those of the WT, although we did not observe the same trend in auxin-induced autonomous seeds (**Fig. S6**).

We then analyzed *bes1-D* and *bzr1-D* mutants, which are constitutive BR signaling mutants. Both alleles result from a single nucleotide change in the coding region which prevents BES1/BZR1 phosphorylation by BIN2, resulting in constitutive accumulation of these transcription factors in the nucleus (Wang et al., 2002; Yin et al., 2002). Unexpectedly, our results showed that 3 DAP seeds of *bes1-D* and *bzr1-D* mutants were smaller than those of WT (**Fig. 3B**). We also observed similar results for autonomous seeds (**Fig. S6**). These findings suggest that, although BR function is required for seed coat formation, excessive BR signaling has a detrimental effect on development. Thus, BRs seem to affect the seed coat development in a dose development manner.

To further verify this, we performed exogenous treatments of WT ovules with epi-brassinolide (epi-BL), a bioactive BR. We treated the unfertilized ovules in a similar manner as we did for the exogenous auxin treatments. To test if exogenous BRs would have an effect on seed coat development, we treated emasculated pistils of Col-0 with different concentrations of epi-BL (100 pm, 1 µM, 10 µM, 100 µM and 150 µM) one day after an exogenous application of 100 µM 2,4-D. Our results revealed that different concentrations of epi-BL had varying effects on seed size: the ovules treated with 100 µM 2,4-D plus 100 pM epi-BL were slightly but significantly larger than those treated with auxin alone, while ovules treated with concentrations equal or above 10 µM of epi-BL were smaller in size compared to the auxin control (**Fig. 3F**). These results support the hypothesis that BRs affect seed growth in a dose-dependent manner, where low concentrations of epi-BL have a positive effect on seed coat development, but concentrations above a certain threshold lead to detrimental effects on seed growth. A similar observation was made when we treated 1 DAP pollinated siliques with Bikinin, a synthetic chemical which activates BR signaling by inhibiting BIN2 (Rybel et al., 2009). We treated the siliques 1 day after pollination with 10 µM, 20 µM, 100 µM and 200 µM of bikinin. The seed size peaked at the 20 µM bikinin treatments, and higher concentrations lead to inhibition of seed coat expansion (**Fig. 3G**). These observations support a dose-dependent effect of BR in seed coat growth, which fits with similar observations made in roots (Vukašinović et al., 2021), in endosperms (Lima et al., 2023) and in pollen tubes (Vogler et al., 2014).

### Seed coat defects in BR mutants are of sporophytic origin

To further verify the origin of the seed coat defects in BR mutants, we compared seed size at 3 DAP in homozygous vs heterozygous mutants. The logic behind this experiment is that if the effect in BR mutants is sporophytic, the heterozygous mutant seeds should behave phenotypically like WT, since the seed coats in heterozygous mutants are diploid and still carry a WT allele. While if the effect in the BR mutant is zygotic, then in a heterozygous condition 25% of seeds will carry mutant embryos and endosperms, and we should see a measurable effect in the phenotype. Along with *bri1* and *det2*, we also used the stronger *dwf4-44* BR biosynthesis mutant for this experiment. The *dwf4-44/-*mutant is extremely dwarf and has severe ovule defects, which prevents its use for reproductive studies in the homozygous state (Lima et al., 2023). But *dwf4-44/+* mutants are phenotypically similar to WT, and produce full seed sets. Our results showed that while seeds of the homozygous *bri1* and *det2* mutants are significantly smaller than WT, as we demonstrated above, the seeds of heterozygous *bri1/+*, *det2/+* and *dwf4-44/+* mutants are indistinguishable from WT (**Fig. 4A-D**). Together with the observation that BR genes are specifically expressed in the sporophytic tissue of seeds (Lima et al., 2023), these results indicate that the small size of BR mutant seeds is due to a seed coat defect.

**Fig. 4.**
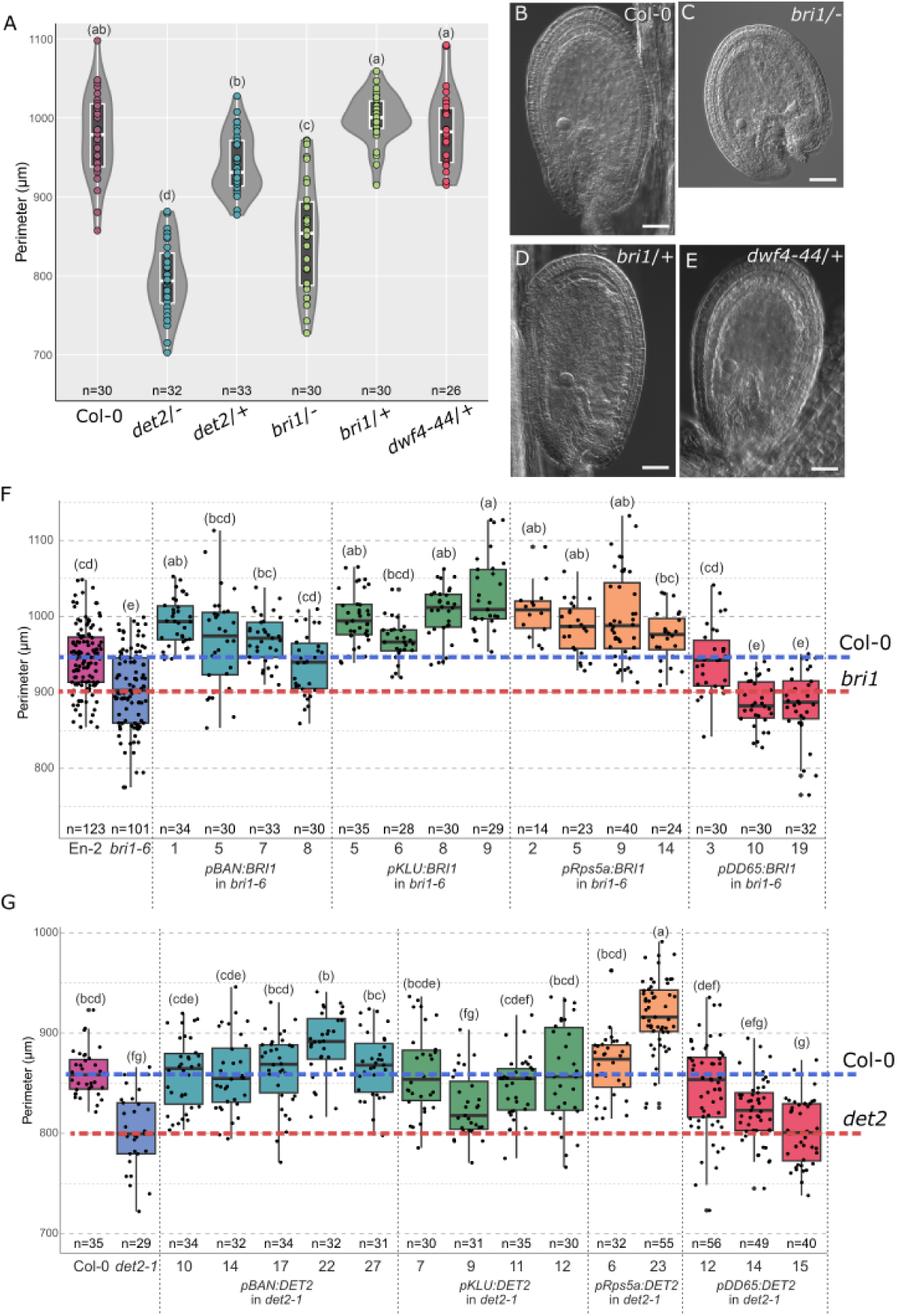
Sporophytic BR activity modulates seed coat growth: (**A**) Seed perimeter of WT, *det2/-, det2/+, bri1/-, bri1/+* and *dwf4-44/+* 3 DAP seeds. Seed morphologies can be seen in **B-E** for WT, *bri1/-*, *bri1*/+ and *dwf4-44*/+, respectively. Scale bars indicate 50 µm. (**F-G**) Perimeter of 3 DAP seeds of *bri1-6* (**F**) and *det2-1* (**G**) and respective complementation lines. All lines are in the *bri1-6* or *det2-1* mutant backgrounds. The phenotypes of the WT and mutants are indicated by the blue and red dashed lines, respectively. The letters indicate statistical significance for p-value <0.05 (Anova).

To further confirm this, we generated tissue-specific rescue constructs to complement the BR mutants. Namely, we complemented the *bri1* and *det2* mutants by expressing the respective genes under the following promoters: *DD65*, specific to the central cell and early endosperm (Steffen et al., 2007); *BANYULS* (*BAN*), specific to the endothelium of the inner integument (Debeaujon et al., 2003); and *KLU*, which is expressed in several integument and seed coat layers (Adamski et al., 2009). Importantly, although *KLU* is expressed in many vegetative tissues, in seeds it is specific to the seed coat at the stage at which we carried out our experiments. Finally, we used the promoter of *Rps5a* (Maruyama et al., 2013), which is constitutively expressed. The *BRI1* and *DET2* coding sequences were cloned under the control of all these promoters and were transformed into *bri1* and *det2* mutants, respectively.

We observed that in the case of *bri1*, several complementation lines showed a rescue of the seed coat growth phenotype. Out of four lines each expressing *BAN::BRI1* or *KLUH::BRI1* in *bri1*, all of them exhibited rescue, i.e., the 3 DAP seed size was restored to WT levels (**Fig. 4F**). This is interesting because *BAN* is only expressed in the innermost layer of the integuments, as compared to *KLU*, which has a broader expression. Native BRI1 is expressed more strongly in the outer integument than in the inner integument (**Fig. S7**) (Lima et al., 2023). This means that restoration of BRI1 expression in one of the integument layers is to some degree sufficient to rescue the *bri1* phenotype. Fitting with the gametophyte-specific expression of *DD65*, two out of three lines expressing *DD65::BRI1* in *bri1* did not show a rescue of the phenotype (**Fig. 4F**). Finally, as expected, all lines expressing *Rps5A:BRI1* showed a rescue of seed size of the *bri1* mutant (**Fig. 4F**).

The same tissue-specific complementation approach was carried out for *det2*. Again, we observed a rescue of the *det2* seed growth defects in eight out of nine lines expressing either *BAN::DET2* or *KLUH::DET2* (**Fig. 4G**). This indicates that restoring BR biosynthesis in the sporophytic tissues is sufficient to rescue the *det2* seed defects. However, it is interesting to point out a particularity about the rescue of *det2* in *BAN::DET2*-expressing lines: endogenous *DET2* is expressed in the outer integument layers (**Fig. S7**) (Lima et al., 2023), but *BAN::DET2* is only expressed in the innermost layer of the integuments (endothelium). Thus, this rescue implies that BR intermediates, produced in inner integuments, can move to outer integument layers, where enzymes that catalyze the next steps of the pathway are located (Lima et al., 2023). Finally, one out of three *DD65::DET2* lines also showed a rescue of the *det2* phenotype (**Fig 4G**). Again, this was surprising given that the *DD65* promoter is specific to the central cell and to the early endosperm, which are symplastically isolated from the seed coat (Stadler et al., 2005). This observation together with the one described above, where the *bri1* phenotype was rescued in one line expressing *DD65:BRI1* might be a sign of non-specific expression of *DD65*. Or alternatively, BR intermediates can cross the endosperm-seed coat barrier and complement the lack of a functional *DET2* in the seed coat of *det2* mutants. In conclusion, our data supports a sporophytic mode of action for seed-produced BRs.

### BR function during seed coat formation is linked to deposition of H3K27me3

Because BR effectors have been shown to recruit ELF6 and REF6 to target loci (Li et al., 2018; Yu et al., 2008), we hypothesized that BR function during seed coat formation could be linked to altered dynamics of H3K27me3. To test if BR effectors and JMJ H3K27me3 are acting in the same pathway during seed formation, we crossed the dominant BR signaling mutant *bzr1-d* to a line ectopically expressing *CaMV35S::ELF6* (Keyzor et al., 2021). However, we observed a high penetrance of aborted and malformed ovules in the double mutant (**Fig. S8**). This was not observed in the single mutants, indicating that indeed BR signaling and H3K27me3 demethylases work in the same pathways during reproductive development. However, this also meant that the double gain-of-function mutants form very few viable seeds and are therefore not useful for functional studies. Thus, as an alternative, we tested if *jmj* mutants are less sensitive to exogenous epi-BL treatments. If BRs act in seed coat development in a manner dependent on H3K27me3 removal, then loss of JMJ function should lead to BR-insensitive seeds. Indeed, the *ref6c elf6* double mutant seeds do not seem to respond to exogenously applied epi-BL (**Fig. S8**).

To further test if BR function in seed coat development is indeed linked to H3K27me3 homeostasis, we focused on the analysis of sporophytic PRC2 mutants, which lack H3K27me3 marks. Loss of PRC2 function in the ovule integuments, such as in a *swn clf*/+ mutant, leads to autonomous seed coat growth (Figueiredo et al., 2016; Roszak and Köhler, 2011). This is because seed coat development pathways are ectopically activated when H3K27me3 is depleted in the ovule integuments. If we hypothesize that BR function during seed coat development is linked to removal of H3K27me3, then *swn clf*/+ mutants should be insensitive to exogenous applications of epi-BL, as they already lack the repressive epigenetic marks. Indeed, we observed little to no effect of exogenous epi-BL applications on *swn/-clf/+* seed size, unlike what happens in the WT (**Fig. 5A**). These results suggest that due to the lack of H3K27me3 marks in *swn/-clf/+* mutants, seed coat growth genes are expressed independently of fertilization and, therefore, BR levels do not affect seed growth.

**Figure 5.**
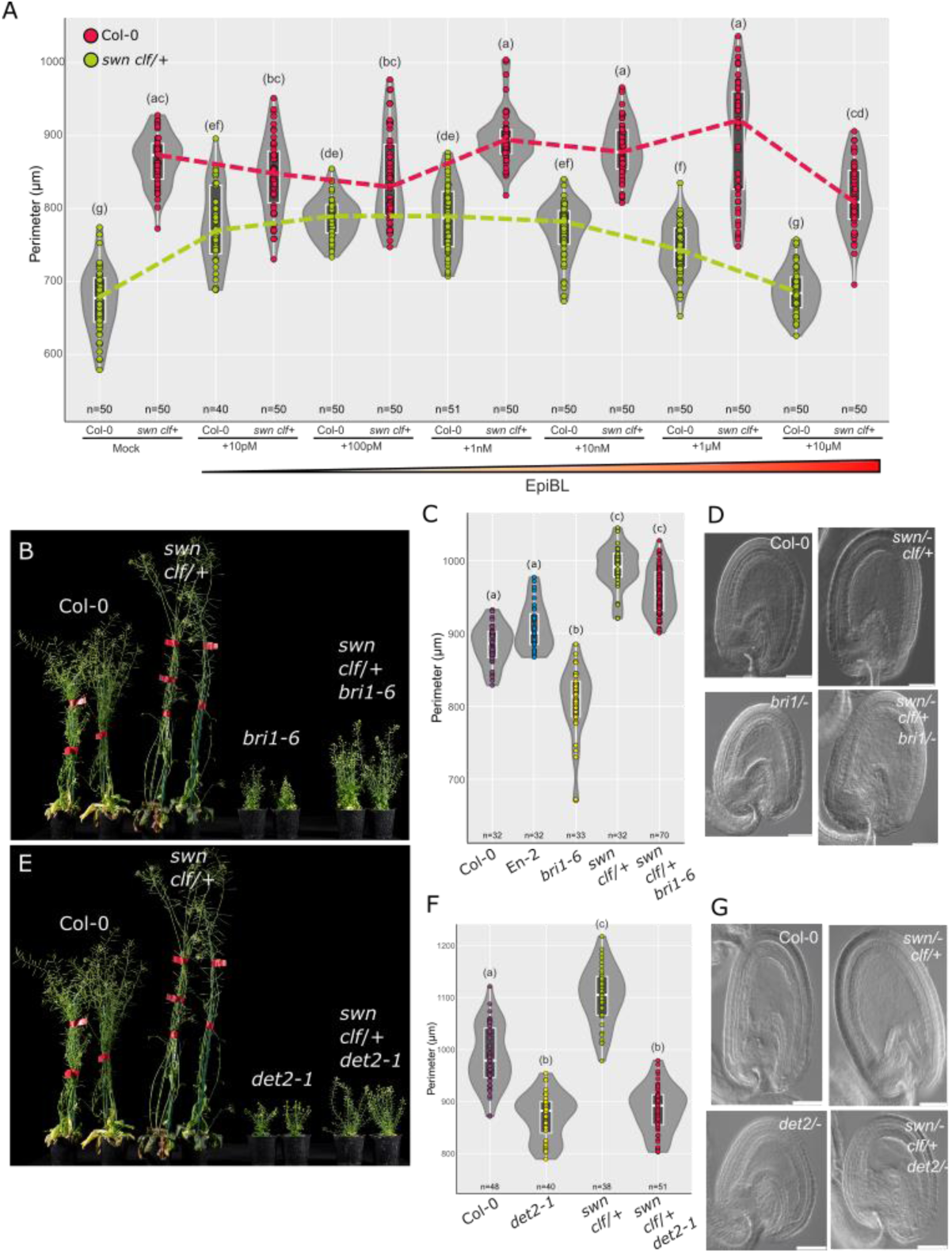
Genetic interactions between PRC2 and BR machinery. (**A**) Seed perimeter of Col-0 (green) and *swn/- clf/+* (red) autonomous seeds at 3 DAT after application of 100 µM 2,4-D and varying concentrations of Epi-BL. (**B**) Vegetative phenotypes of WT, *swn/- clf/+*, *bri1-6* and corresponding triple mutant. (**C**) Seed perimeter at 3 DAP of WT, *swn/- clf/+*, *bri1-6* and corresponding triple mutant. (**D**) Autonomous seed morphologies of WT, *swn/- clf/+*, *bri1-6* and *swn/- clf/+*, *bri1-6*. Scale bars indicate 50 µm. (**E**) Vegetative phenotypes of WT, *swn/- clf/+*, *det2-1* and corresponding triple mutant. (**F**) Seed perimeter at 3 DAP of WT, *swn/- clf/+*, *det2-1* and corresponding triple mutant. (**G**) Autonomous seed morphologies of WT, *swn/- clf/+*, *bri1-6* and *swn/- clf/+*, *det2-1*. Scale bars indicate 50 µm. The letters in (**A,C,F**) indicate statistical significance for p-value <0.05 (Anova).

Moreover, if our hypothesis is true that BR function in the seed coat is necessary for the efficient removal of H3K27me3 marks by JMJs, then we expect that the seed coat defects of BR mutants are alleviated by loss of PRC2, because those epigenetic marks are not deposited to start with. Thus, we crossed the sporophytic PRC2 mutant *swn/-clf/+* with BR loss-of-function mutants. Interestingly, in the case of *swn/-clf/+ bri1/-* we observed that the triple mutant plants displayed some rescued plant morphologies compared to the *bri1* plants. The triple mutant had larger and more leaves, grew taller (**Fig. 5B**), and flowered later but also for a longer time than *bri1* plants. Importantly, we also observed a similar rescue of the *bri1* growth defects in sexual and autonomous seeds (**Fig. 5C-D and S8**). In fact, seeds of *swn/-clf/+ bri1/-* were of the same size of those of *swn/-clf/+* mutants, resulting in a complete rescue of the growth phenotype and demonstrating that loss of PRC2 is epistatic to the loss of BR signaling via BRI1. Additionally, we observed that the unfertilized ovules of *swn/-clf/+ bri1/-* were of the same size as those of WT and *bri1/-*, but smaller than those of *swn/- clf/+* (**Fig. S8**). This suggests that lack of BRs to some degree may repress the development of autonomous seed coats in *swn/- clf/+*, potentially because of inefficient removal of residual H3K27me3.

Next, we did the same experiment but using the *det2* BR biosynthesis mutant. Surprisingly, the outcome was different from that obtained for *swn*/- *clf*/+ *bri1*/-. While we did observe some rescued plant morphologies in *swn/- clf/+ det2/-*, this was not as striking as for *swn/- clf/+ bri1/-* (**Fig. 5B,E**).

Moreover, unlike for *bri1*/-, loss of PRC2 function did not rescue the seed growth phenotype of *det2*/-, either in fertilized or auxin-induced autonomous seeds (**Fig. 5F-G and S8**). This suggests that the epigenetic control of seed coat development through BRs might function through multiple pathways, some independent of the main receptor BRI1. To further validate this, we crossed *swn/- clf/+* to another BR biosynthesis mutant, *dwarf4-102/+* (*dwf4*). Homozygous mutants for *dwf4-102/-* are severely dwarf and cannot be used for reproductive studies. However, heterozygous *dwf4-102/+* mutants are haplo-insuficient and their 3 DAT seeds are smaller than those of the WT (**Fig. S8**). Importantly, similar to what we observed for *det2*, loss of sporophytic PRC2 did not result in a rescue of the *dwf4-102*/+ phenotype (**Fig. S8**). This suggests that, unlike what happens for BR signaling via BRI1, loss of BR biosynthesis is epistatic to loss of PRC2 and, thus, of H3K27me3.

Our genetic analysis supports the hypothesis that BR-mediated seed coat growth, via the main receptor BRI1, works through H3K27me3 removal. If this is true, *bri1* mutants should be impaired in the removal of this epigenetic mark. To test if this is the case, we profiled H3K27me3 in whole fruits of *bri1-6* and the respective WT (En-2) at 3 DAP (**Fig. 6 and S9**). In both cases the WT was used as a pollen donor, so that any effects that we detect are attributable to H3K27 profiles in the sporophyte and not in the fertilization products, embryo and endosperm. We divided genes in five clusters, according to their H3K27me3 enrichment (**Table S2**). Clusters 1 to 3 contain genes enriched in H3K27me3 in the coding region, or in the promoter and terminator regions. While Clusters 4 and 5 contain genes with little to no H3K27me3 signal in their vicinity. A list of genes specific to cluster 1, which are labeled the strongest with H3K27me3, can be found in **Table S3**. As hypothesized, *bri1* shows a global enrichment in H3K27me3, when compared to the WT (**Fig. 6A and S9**). For instance, while in WT we detected 10,409 genes in the top three clusters, this number rose to 14,447 genes in *bri1*. This signifies a 39% increase in the number of genes labeled with H3K27me3 when BRI1 is not present. We also checked if genes placed in Cluster 1 for *bri1* (strongly labeled) were demethylated in WT, and thus placed in lower-ranking Clusters. Indeed, 867 genes can be found in Cluster 1 of *bri1*, but not present in the same cluster in WT, and many such genes are described as expressed in seeds (**Fig. 6B, Table S3**). Additionally, 507, 391 and 21 genes that are found in Cluster 1 of *bri1* are in turn present in the WT Clusters 2, 3 and 4, respectively (**Fig. 6C-E**). This indicates that genes that are normally demethylated in a WT condition are ectopically methylated in *bri1* (example in **Fig. 6F**). We then took these genes which are strongly methylated in *bri1* but lose this methylated status in WT, and did a gene ontology (GO) term enrichment analysis (**Fig. 6G** and **Table S4**). Interestingly, we identified GO terms related to secondary metabolism, namely phenylpropanoid biosynthesis, which is a hallmark of seed coat formation (Debeaujon et al., 2003). Finally, to test if H3K27me3 hypermethylation could be linked to JMJ activity, we tested for the presence of REF6 *cis-*binding motifs, CTCTGYTY, in genes belonging to *bri1* cluster 1. Indeed, genes hypermethylated in *bri1* are enriched in REF6 binding motifs with their promoter regions, when compared to the control (**Fig. 6H**). Thus, this H3K27me3 profiling supports the hypothesis that signalling via BRI1 is required for H3K27me3 removal, likely via the activity of JMJ histone demethylases, and is likely required for the activation of genes involved in seed coat formation.

**Figure 6.**
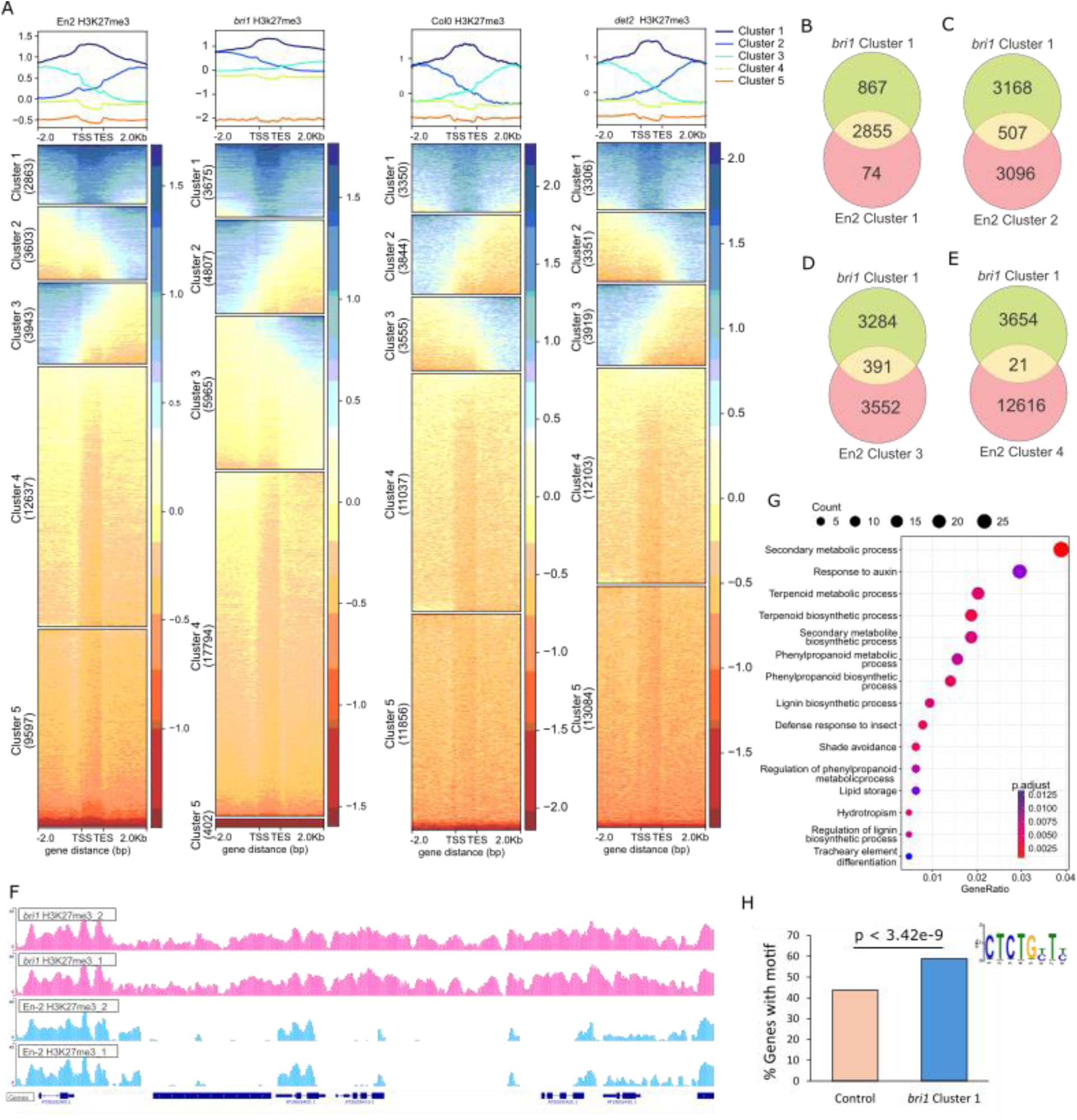
Loss of BRI1 leads to H3K27 hypermethylation. **A** Clustering analysis of H3K27me3 profiles normalized to H3. The metagene plots on top indicate the average enrichment of H3K27me3 over gene regions for each of the Clusters indicated below. The colors indicate relative enrichment. TSS - transcription start site, TES - transcription end site. One replicate is shown for each genotype. Duplicates can be found in **Fig. S9** and **S10**. **B-E** Overlap of genes present in *bri1* Cluster 1 with those present in WT Clusters 2 (**C**), 3 (**D**) and 4 (**E**). (**F**) H3K27me3 profiles illustrating H3K27me3 hypermethylation in *bri1*, when compared to the respective En-2 WT. Two biological replicates per genotype are shown. (**G**) GO enrichment analysis for genes enriched in H3K27me3 in *bri1*, compared to En-2 WT. (**H**) Enrichment of CTCTGYTY motifs in the 3 kb promoter region of genes in *bri1* cluster 1, compared to a control sample.

We then repeated the CUT&TAG H3K27me3 analysis using the *det2* mutant and respective WT (Col-0). Again, in both cases WT was used as a pollen donor. Unexpectedly, unlike we observed for *bri1*, there were no obvious signs of H3K27me3 hypermethylation in *det2* (**Fig. 6A and S10**). The number of genes present in each *det2* cluster was comparable to those detected in the WT samples (**Fig. 6A and Table S2**). Moreover, there was a strong overlap between the genes within all WT and *det2* clusters, which was not the case for *bri1* (**Fig. S11**). This suggests that BR biosynthesis, unlike signalling via BRI1, is not required for H3K27me3 removal.

The observation that loss of signalling via BRI1 leads to H3K27 hypermethylation, but that loss of BR biosynthesis does not, suggests that BRI1-mediated H3K27me3 removal works in a BR-independent manner. This fits with the genetic data of **Fig. 5B-G**, where *bri1* mutant phenotypes can be rescued by loss of H3K27me3, but this is not the case for *det2* phenotypes. This led us to hypothesize that BRI1-mediated H3K27me3 removal may create a permissive environment for BR perception and responses. Thus, we tested whether the BR-insensitivity of BRI1 mutants is indeed due to the lack of BR perception by BRI1, or to the absence of a permissive environment for BR to act. To do this, we tested the inhibition of root elongation by exogenous epi-BL (**Fig. S12**). Both WTs tested, Col-0 and En-2 showed a stark inhibition of root elongation when exposed to 100 nM epi-BL. The same was true for the PRC2 mutant *swn clf*/+. In contrast, *bri1*-*6* roots were insensitive to epi-BL and even elongated further when exposed to the chemical. And the same phenotype was true for the triple mutant *bri1-6 swn clf*/+. This means that BRI1 is necessary for BR perception, even when H3K27me3 is depleted. Thus, although loss of H3K27me3 is epistatic to the seed growth phenotypes of *bri1-6* (**Fig. 5**), loss of BRI1 is epistatic to H3K27me3 hypomethylation when it comes to root elongation (**Fig. S12**). Therefore, this suggests that although some *bri1* phenotypes are indeed due to its ability to transduce BR signals, BRI1 also functions in a BR-independent manner, when it comes to H3K27me3 removal.

### BRI1-independent pathways also control seed coat formation

Although our data points to BRI1 having BR-independent functions in mediating H3K27me3 removal, loss of BR biosynthesis does result in impaired seed coat development (**Fig. 3B,E**), but this likely happens in a manner independent of H3K27me3 removal and, thus, independently of BRI1 (**Fig. 5B-G**). Alternative pathways through which BR may mediate seed coat growth could be via BRI1-LIKE receptors (BRLs). There are three BRL receptors in Arabidopsis, of which only two bind bioactive BRs, BRL1 and BRL3 (Caño-Delgado et al., 2004). Based on published gene expression data, only *BRL3* is expressed in the seed coat (**Fig. S13**). To confirm this, we generated a *BRL3::NLS:tdtomato* reporter line and observed that *BRL3* is strongly expressed in the chalazal seed coat (**Fig. 7A**). Then, to investigate if BR signaling through BRL3 could be involved in seed coat formation, we analyzed 3 DAT and 3 DAP seeds in a *brl3/-* mutant. Indeed, both autonomous and fertilized seeds were significantly smaller in the mutant than in the WT (**Fig. 7B** and **S13**), despite *brl3/-* mutant plants looking phenotypically similar to WT.

**Figure 7.**
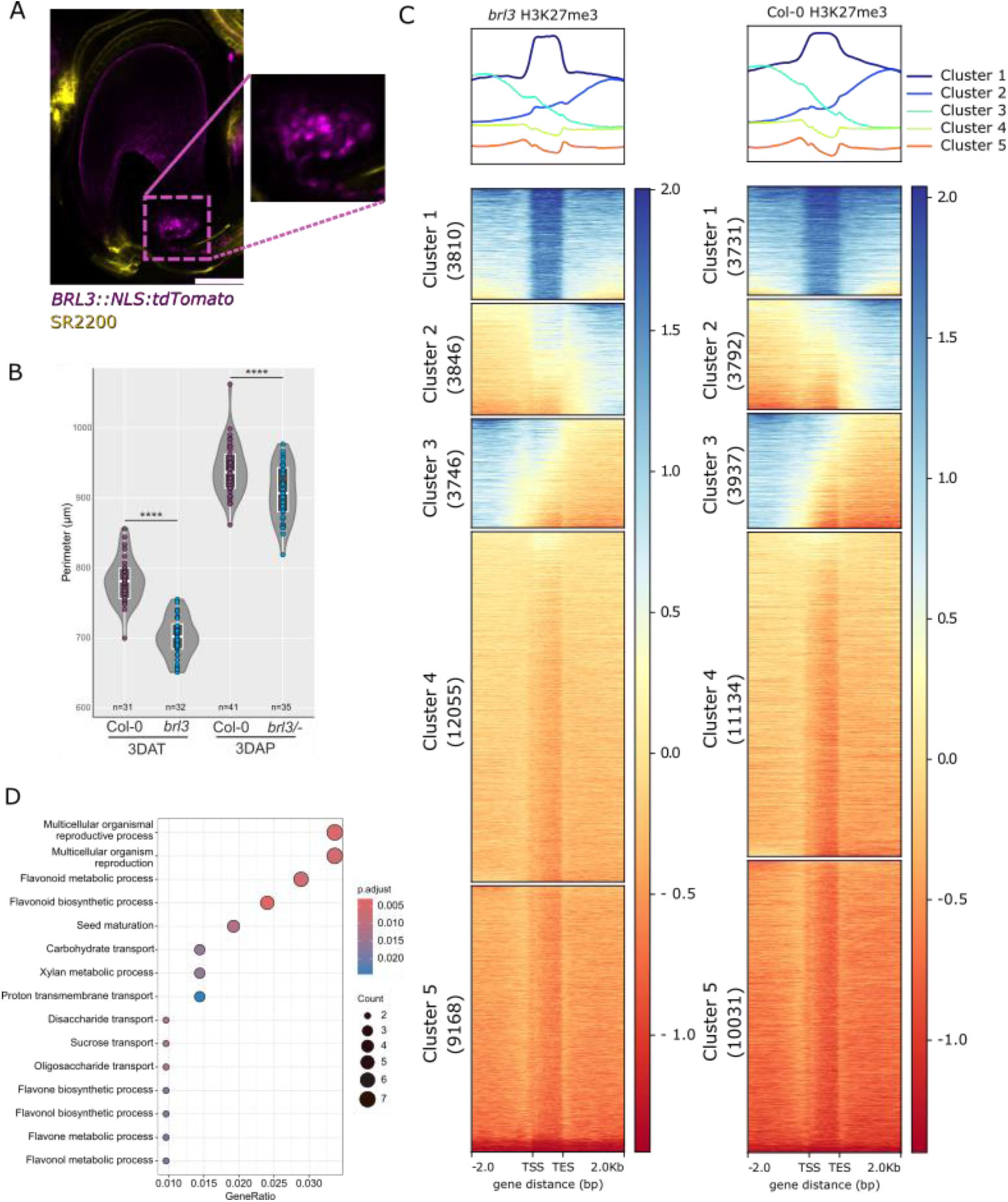
BRL3 regulates seed coat formation independently of H3K27me3. (**A**) Relative expression of BRI-LIKE genes in seed coat, at different stages of seed development, as determined by (Belmonte et al., 2013). (**B**) Expression of a *BRL3::NSL:tdtomato* reporter in a seed at 2 DAP. Yellow is a counterstain with SR2200. Scale bar indicates 50 µm. (**B**) Seed perimeter of Col-0 and *brl3* 3 DAP fertilized and 3 DAT autonomous seeds. **** indicates p-value <0.0001. (Anova). (**C**) Clustering analysis of H3K27me3 in *brl3* and in Col-0 normalized to H3. The metagene plots on top indicate the average enrichment of H3K27me3 over gene regions for each of the Clusters indicated below. The colors indicate relative enrichment. TSS - transcription start site, TES - transcription end site. One replicate is shown for each genotype. Duplicates can be found in **Fig. S14**. (**D**) GO enrichment analysis for genes enriched in H3K27me3 in *brl3*, compared to Col-0 WT.

Next, to test if the regulation of seed coat formation by BRL3 is also related to the removal of H3K27me3 marks, we crossed the *swn/+ clf*/- mutant with *brl3* to examine whether the *swn/- clf/+ brl3/-* triple mutants exhibits any rescue in seed size as observed for *swn/- clf/+ bri1/-* (**Fig. 5C**). However, we only observed a slight rescue of the *brl3*/- seed phenotype in *swn/- clf/+ brl3/-* for both sexual and autonomous seeds (**Fig. S13**). Although the *swn/- clf/+ brl3/-* triple mutant seeds were bigger than those of the *brl3/-* single mutant, they were only as large as WT seeds. And nowhere near those of the *swn/- clf/+* double mutant, which is what we observed in the case of *bri1* (**Fig. 5C**). This suggests that the *brl3* seed phenotypes are only partly related to H3K27me3 removal, and that BRL3 may work directly in BR signal transduction during seed coat formation. To test this, we profiled H3K27me3 on *brl3* and Col-0 siliques. Again, in both cases WT pollen was used. As hypothesized, *brl3* H3K27me3 profiles are only slightly different from those of the WT (**Fig. 7C** and **Table S2**). This more resembles those profiles obtained from *det2*, than those of *bri1* (**Fig. 6A**). Nevertheless, we did detect some genes which showed H3K27me3 hypermethylation in *brl3*, when compared to the WT. We did a GO enrichment analysis with those genes and found processes that are also related to seed coat formation (**Fig. 7D** and **Table S4**). Such terms include flavonoid metabolism and sugar transport. Interestingly, several of those genes are known regulators of sugar import during seed formation, and they form an interaction network (**Fig. S14**). Thus, BRL3 does seem to be required for H3K27me3 removal in a small subset of genes, which are required for seed coat development. This function of BRL3 is likely independent of BR. But because the *brl3* phenotypes are not fully rescued by H3K27me3 depletion, BRL3 does seem to work in both BR-dependent and -independent manners during seed coat formation.

We thus propose a model where BR signaling and H3K27me3 removal by JMJ histone demethylases work in a coordinated manner to allow seed coat development. In it, BR signaling through the main receptor BRI1 is necessary for seed coat formation in a manner dependent on H3K27me3 removal, likely through recruitment of ELF6 and REF6 by BR effectors. While signaling through BRL3 is also necessary for seed coat growth, but in a manner mostly independent of H3K27me3 removal (**Fig. 8**).

**Fig. 8.**
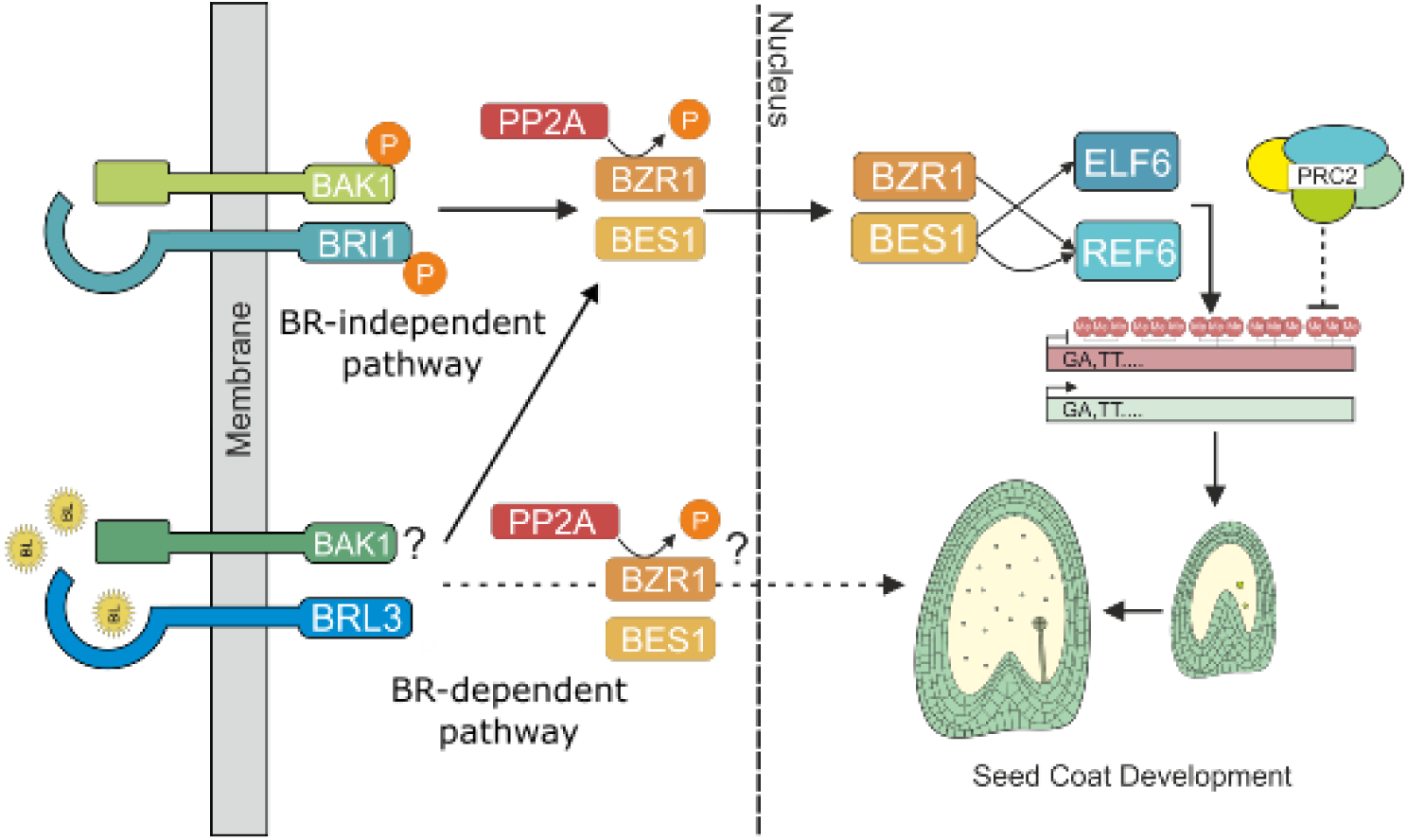
Working model for the regulation of seed coat formation by BR-related pathways. BRI1-mediated signaling is necessary for removal of H3K27me3 marks from the integuments, priming this tissue for seed coat development. This process is independent of BRs and is likely done in coordination with JMJ H3K27me3 demethylases. BRL3-mediated signaling pathways are also necessary for seed coat growth. In this case, these pathways are also involved in mediating H3K27me3 removal, but also likely work in a BR-dependent fashion.

## Discussion

Deposition of H3K27me3 prevents seed coat development prior to fertilization and these repressive marks must be removed in order for the seed coat to form properly (Figueiredo et al., 2016; Roszak and Köhler, 2011). Our findings support the hypothesis that removal of H3K27me3 is most likely carried out by members of the JMJ family of histone demethylases. Several pieces of evidence support this: first, REF6 and ELF6 are expressed in the integuments and in seed coats. Second, high order *jmj* mutants show seed coat formation defects, including slower relative growth and delayed accumulation of proanthocyanidins, a hallmark of seed coat formation in Arabidopsis (Debeaujon et al., 2003). This phenotype contrasts to that observed in mutants for PRC2, which accumulate proanthocyanidins even without fertilization (Figueiredo et al., 2016), and thus links H3K27me3 removal to seed coat formation.

Moreover, we observed additional reproductive phenotypes in the *elf6 ref6c jmj13* triple mutant, including decreased ovule viability and pollen function, leading to fewer fertilized seeds. This decrease in seed set in the triple mutant siliques may result in more resources being directed towards the remaining seeds, potentially mitigating the putative seed coat defect phenotype found in this triple mutant. Such inverse correlations between seed set and size are well documented in the literature (Jofuku et al., 2005; Ohto et al., 2005). However, we did observe that even single *jmj* mutants produced seeds at maturity that were larger than their WT counterparts, even if their seed set was not much compromised. Our data suggests that zygotic effects from the embryo and the endosperm contribute to this phenotype. Indeed, JMJ function had already been demonstrated in endosperm development: REF6 was shown to remove H3K27me3 marks from the maternal alleles of the endosperm, activating them during germination (Sato et al., 2021). However, we cannot rule out a dual-role of JMJ function in the seed coat, first being necessary for seed coat initiation, by removal of H3K27me3 marks from the integuments, and later working to repress seed expansion, potentially by targeting different loci at different stages of development. This would fit with REF6 targeting different sets of genes in different stages of seed development.

Given the documented interactions between BR effectors and JMJ demethylases, namely ELF6 and REF6 (Li et al., 2018; Yu et al., 2008), we then assessed the potential role of these steroid hormones in regulating seed coat formation. Based on previously published datasets (Belmonte et al., 2013), BR related genes were predicted to be expressed in seed coats, similar to what happens with *JMJ* genes. We thus hypothesized that BR signaling would be required for JMJ function during seed coat development, allowing for H3K27me3 to be removed from the integuments. In particular, the expression of the BRI1 receptor in the integuments and seed coat, as well as the strong seed coat defects of *bri1*, confirms that BR signaling is required for proper seed coat formation. Both BRI1 and BZR1 were previously shown to be expressed in the integuments during ovule development (Jia et al., 2020). This suggests that BRs are also necessary during ovule development before fertilization. Whether this is via the regulation of H3K27me3 homeostasis or not, is unknown, although it would fit with our observations that lines ectopically expressing *CaMV35S::ELF6* and *bzr1-D* have severe ovule developmental defects.

Consistent with a role for BR in seed coat formation, BR mutants exhibited smaller seed sizes compared to the WT. Surprisingly, constitutive BR signaling mutants also exhibited smaller seed sizes compared to the WT. These findings suggest that excessive BR signaling has a detrimental effect on seed coat development, indicating a dose-dependent response of BRs in seed coat growth. Similar observations have been made in other organs (Lima et al., 2023; Vogler et al., 2014; Vukašinović et al., 2021). Interestingly, it was previously shown that BES1 is ectopically dephosphorylated in *dwf4-5D,* suggesting increased BR signaling in this gain-of-function mutant (Kim et al., 2013). The reason why *bes1-D* has a negative effect on seed size but *dwf4-5D,* which also presumably has an increased activity of BES1, has a positive effect is still an open question. This dose-dependent mode of action for BRs on seed coat formation was confirmed with exogenous treatments of ovules with epi-BL and with bikinin. For both chemicals, low concentrations had a positive effect on seed coat growth, which was negated at higher concentrations. While the reason for this remains unknown, it is possible that extreme BR levels could result in early or excessive recruitment of ELF6/REF6, resulting in ectopic H3K27me3 demethylation.

Interestingly, BRs have been previously proposed to regulate seed growth and shape in Arabidopsis, via the direct regulation of the endosperm-specific genes *SHB1*, *IKU1*, *MINI3*, and *IKU2* by BZR1 (Jiang et al., 2013). These genes were thus proposed to regulate seed size downstream of DET2 and BZR1 (Jiang et al., 2013; Jiang and Lin, 2013). However, *MINI3* seems to be also expressed in the sporophytic tissues of the seed (Kang et al., 2013). Alternatively, BZR1 could directly regulate *IKU2* expression (Jiang et al., 2013; Jiang and Lin, 2013), which would require BR signaling to be active in the endosperm. In contrast, our data points to a sporophytic effect on seed size and, furthermore, our reporter analysis places all BR effectors in the seed coat and not the endosperm (Lima et al., 2023). However, we cannot rule out that BR has zygotic effects on seed growth at later stages than the ones we assessed in this study.

We went further to test whether the seed coat defects of BR mutants were indeed linked to poor removal of H3K27me3. Indeed, loss of sporophytic PRC2 alleviates the seed coat defects observed in *bri1*. Notably, *swn/- clf/+ bri1*/- exhibited rescued plant morphologies, including increased leaf size and plant height. This means that many reported phenotypes observed in BR mutants are likely due to altered H3K27me3 homeostasis. Importantly, loss of H3K27me3 in the seed coat is epistatic to the loss of BR signaling via BRI1. This was further confirmed by the observations that the PRC2 mutant was insensitive to exogenous BR applications. A surprising observation came from our analysis of the *swn/- clf/+ det2/-* triple mutant, whose outcome was the opposite of *swn/- clf/+ bri1/-*. Although some rescue of plant morphologies was observed in the triple mutant, it was not as significant as that observed for *bri1*. Moreover, the seed size of *swn/- clf/+ det2/-* remained the same as for the single *det2/-* mutant, indicating that loss of BR biosynthesis is epistatic to loss of PRC2. This suggested that the development of seed coat through BRs may involve multiple pathways, either dependent or independent of the main receptor BRI1. We hypothesized that such an alternative pathway could be under the control of BRLs (Caño-Delgado et al., 2004). Indeed, *brl3* mutants have smaller seeds compared to the WT, and *BRL3* is specifically expressed in the chalazal seed coat. This indicates that BR signaling through BRL3 is involved in seed coat formation. Importantly, the seed coat defects of *brl3* are not rescued by loss of PRC2, unlike what happens for *bri1*. These observations were supported by H3K27me3 profiling in these three mutants. While loss of BRI1 leads to H3K27me3 hypermethylation in many loci, the same is not true for loss of DET2 or of BRL3. This supports the hypothesis of a BRI1-independent pathway which affects the seed coat growth directly, and not through modulation of H3K27me3 levels. This observation was very interesting because BRI1 and BRL3 are homologous proteins with presumably similar functions. BRL3 was shown to complement the *bri-301* mutant when expressed under the endogenous *BRI1* promoter (Caño-Delgado et al., 2004). It is not known yet if both BRI1 and BRL3 activate the same or different components of BR signaling, but our results show that they might be active in different pathways. It was previously hypothesized that BES1, which is activated when BRI1 senses BRs, itself seems to act as an activator of BRL3 at low levels of BR, whereas at higher levels, BES1 acts as a repressor of BRL3 in roots (Salazar-Henao et al., 2016).

Regarding the BR-independent regulation of seed coat formation by BRI1, which is linked to poor H3K27me3 removal, it remains to be determined how these pathways are transduced. Indeed, BRI1 has been reported to have functions independent of BRs (Holzwart et al., 2020). Moreover BRI1 internalization can occur independently of its binding to the hormone (Neubus Claus et al., 2023). In the future it will be interesting to elucidate how BRI1 mediates H3K27me3 removal, and how this pathway can be uncoupled from BRs. The same is true to how BRL3 and BRI1 both signal to regulate the same developmental process, but via independent molecular pathways.

Overall, our study provides evidence for the co-involvement of JMJ histone demethylases and of BR-related signaling in regulating seed coat development in Arabidopsis via modulation of H3K27me3 removal. The findings suggest that BR signaling is crucial for seed coat growth and that the interplay between BRs, PRC2-mediated epigenetic control, and BRL receptors contributes to the intricate regulatory network underlying seed coat formation. We thus propose that: 1) BRI1-mediated BR signaling is necessary for H3K27me3 removal from the integuments, allowing for seed coat formation, and 2) BRI1-independent BR signaling is also necessary for seed coat growth, but in a manner mostly independent of H3K27me3 removal.

## Materials and Methods

### Plant material and Growth Conditions

The lines used in this study are *elf6-3* (SALK_074694), *elf6-4* (SAIL371D8), *ref6-1* (SALK_001018), *jmj13* (GABI-Kat113B06), *ref6c, elf6-3ref6c, elf6 ref6c jmj13 (ERJ)* and *REF6::REF6:GFP* (Yan et al., 2018)*, bri1-6* (Noguchi et al., 1999)*, det2* (Chory et al., 1991)*, bzr1- D* (Wang et al., 2002)*, dwf4-5D* (Kim et al., 2013)*, swn-3* (Chanvivattana et al., 2004) *clf-9* (Goodrich et al., 1997) (used as *swn/- clf/+*)*, bri1-301* (Xu et al., 2008), *BRI1OX* (Friedrichsen et al., 2000), *bes1-D* (Yin et al., 2002)*, cpd91* (SALK_078291), *dwf4-102* (SALK_020761), *dwf5-7* (SALK_127066), *brl3* (SALK_006024), *br6ox1* (SALK_148384) *br6ox2* (SALK_056270), *ROT3::NLS:GFP, DWF4::NLS:GFP*, *BR6OX1::NLS:GFP, BR6OX2::NLS:GFP, BES1::BES1:GFP* (Vukašinović et al., 2021), *BRI1::BRI1:GFP* (Sun et al., 2020), *CYP90A1p:NLS:3xEGFP* (*CPD*) (Vogler et al., 2014), *JMJ13::JMJ13:GFP* (Keyzor et al., 2021), *REF6::REF6*, *TPS1::REF6* and *EPR1::REF6* (Sato et al., 2021). The primer sequences for genotyping mutants lines can be found in **Table S1**.

Seeds were sterilized with 5% commercial bleach with 0.01% Triton X100 for 5 mins followed by 3 times washing with 99.6% ethanol. The sterile seeds were plated onto ½ MS-medium supplemented with 1% sucrose. The plates were kept at 4°C for 48h in the dark for stratification. Plates were then transferred to a growth chamber (16 h light/8 h dark; 50 μmol.s^−1^.m^−2^; 22°C). After 10 days, the seedlings were transferred to soil and grown in a growth chamber (16 hr light/8 hr dark; 150 μmol.s^−1^.m^−2^; 21/20°C; 70% humidity).

### Physiological assays

The hormone treatments used contained 0.1% of ethanol, 0.01% Silwett L-77, and 100 μM of 2,4-Dichlorophenoxyacetic acid (2,4-D). To ensure accuracy in the results, a mock control group was also included in all experiments. Two days prior to anthesis, the flowers were emasculated and two days later were treated with 2,4-D or mock solutions or pollinated. At the designated time intervals, usually three days after treatment (3DAT) or pollination (3DAP), the treated pistils were collected and prepared for microscopy examination.

For clearing of ovules and seeds the whole pistils/siliques were fixed with EtOH:acetic acid (9:1), washed for 10 min in 90% EtOH, 10 min in 70% EtOH and cleared overnight in chloral hydrate solution (66.7% chloral hydrate (w/w), 8.3% glycerol (w/w)). The ovules/seeds were observed under differential interference contrast (DIC) optics using a Leica DM2500 microscope (Leica Microsystems). The DMACA staining was done on 1 DAP, 2 DAP, 3 DAP seeds and 3 DAT autonomous seeds in 2% (w/v) DMACA (p-dimethylaminocinnamaldehyde) in (1:1) 6 N HCl:96% EtOH. The emasculated pistils were incubated in this solution for 30 min and then the ovules/seeds were dissected out and mounted on a microscope slide. Images were recorded using a Leica DM2500 microscope. **Fig. S3** shows the criteria for different levels of DMACA staining. Seed perimeter and area were measured from DIC images using Fiji software. Plots and statistical analysis were done in RStudio.

For fluorescence analysis seeds were mounted in water with 0.1 mg/mL propidium iodide (PI). Samples were analyzed under confocal microscopy on Leica Stellaris 8 Dive with the following settings (in nm; excitation-ex and emission-em): GFP – ex 488, em 500–530; PI – ex 488/514, em 635–719, EYFP (VENUS) – ex 514, em 527. Images were acquired, analyzed and exported using LASX software.

For root growth experiments, the seeds were plated on ½ MS plates with or without 100 nM epi-BL. After one week of growth, the plates were imaged using a Keyence Digital Microscope VHX- 6000. Root length was measured using SmartRoot (Lobet et al., 2011). The images were converted to grayscale 8-bit, negated, and converted to black and white with a 60% threshold. Between 60 and 120 roots were measured for each genotype.

### Cloning and generation of transgenic plants

To clone the construct *JMJ13::GFP*, 2300 bps of the *JMJ13* promoter were amplified from Col-0 genomic DNA. The amplified sequence was purified from the gel and was recombined into the donor vector (pDONR221) using BP Gateway cloning according to the manufacturer’s instructions (Fisher Scientific). The donor vector was sequenced and the insert recombined into the destination vector pB7FWG.0 using LR Gateway cloning.

To clone *KLUH:BAS1* and *BAN:BAS1*, the *BAS1* coding region was amplified from Col-0 cDNA. The amplified fragments were purified from the gel and were transferred into donor vector pDONR221 using BP Gateway cloning according to the manufacturer’s instructions (Fisher Scientific). The donor vector was sequenced to confirm the correct sequence. The donor vector carrying the *BAS1* gene was then recombined into two modified pB7WG2 vectors (VIB, Ghent), where the *CaMV35S* promoter had been replaced with either 4100 bp of the *KLUH* promoter or with 355 bp of the *BANYULS* (*BAN*) promoter.

To clone the constructs for complementation of *bri1* and *det2* mutants, the genomic regions of *BRI1* and *DET2* were amplified from Col-0 genomic DNA. The amplified PCR fragments were purified from the gel and were transferred into donor vector pDONR221 using BP Gateway cloning according to the manufacturer’s instructions (Fisher Scientific). Both donor vectors were sequenced and recombined into four modified pB7WG2 vectors using LR Gateway cloning technology. In these destination vectors the *CaMV35S* promoter was replaced with the promoters of the following genes (length of the promoter region indicated in brackets): *DD65* for central cell and early endosperm specific expression (1277 bp), resulting in *DD65:BRI1* and *DD65::DET2* constructs; *BAN* for expression in the endothelium layer of the seed coat (355 bp), resulting in *BAN:BRI1* and *BAN::DET2* constructs; *KLUH* as stronger seed coat specific construct (4100 bp), yielding *KLUH:BRI1* and *KLU::DET2*; and finally *Rps5a* as a constitutively expressed promoter (1613 bp), resulting in constructs *Rps5a:BRI1* and *Rps5a:DET2*.

To clone the constructs for expression analyses of *REF6* and *ELF6*, the pB7WG vector was digested with *Eco*32I and ligated to remove the cassette of the LR reaction. The *GUSplus* reporter gene (Broothaerts et al., 2005) was inserted into *Pst*I and *Bcu*I sites, and the *REF6* and *ELF6* promoter regions (3116 bp and 3002 bp, respectively) were inserted into the *Kpn*I and *Xba*I sites of the vector using the In-Fusion HD Cloning Kit (TaKaRa).

To clone the BRL3 reporter, its promoter was amplified and cloned as a blunt fragment into pJET1.2 (Thermo), and then used to substitute the CaMV35 promoter in pK7WG2 (VIB, Ghent) as a SpeI-SacI fragment. Finally, an NLS:tdTomato cassette was recombined into that vector via LR Gateway cloning. The ENTRY vector was pEN-L1-NTdTomato-St-L2,0 (VIB, Ghent).

The primer sequences used for cloning can be found in **Table S1**. Multiple independent copies of each construct were transformed into *Agrobacterium tumefaciens* GV3101, and then into *Arabidopsis thaliana* plants using floral dip (Clough and Bent, 1998). The transformants were selected on ½ MS-medium supplemented with 1% sucrose and the appropriate selection agent.

### REF6 target prediction

Genes were selected that bear REF6 binding sites, as determined by (Cui et al., 2016), and that were expressed during seed development, as determined by (Belmonte et al., 2013). Only genes bearing four or more CTCTGYTY motifs (N>=4, CTCTGYTY) were included in the analysis. Out of 406 genes meeting this criterion, 326 were expressed in seeds and were used to generate the heatmap of **Fig. S5**. The heatmap was clustered by row with an average Linkage method and Pearson distance measurement method.

To test for the enrichment of REF6 binding sites in genes belonging to the *bri1* cluster, the 3.0 kb regions upstream of genes within that cluster were tested for the presence of CTCTGYTY motifs using SEA (Bailey and Grant, 2021). Control sequences are shuffled preserving 3-mer frequencies. Statistical significance was testing using a Fisher Exact Test.

### Profiling of H3K27me3 using CUT&TAG

The *bri1* x En2, *det2* x Col-0, and *brl3* x Col-0 crosses, along with their respective wild-type (WT) controls, were performed using plants grown in standard growth conditions, as described above. Samples were collected at 3 DAP and snap-frozen in liquid nitrogen for subsequent analysis. Nuclei were extracted from the frozen whole siliques using a GentleMACS dissociator (Miltenyi Biotec, cat. no. 130-093-235), following the protocol established by (Moreno-Romero et al., 2017)

The CUT&TAG protocol was performed as described (Del Toro-de León et al., 2024). Briefly, was performed on each sample from mutant crosses and their corresponding WT controls using the CUTANA pAG-Tn5 kit. Primary antibodies against H3K27me3 (Cell Signaling Technology, 9733T) and H3 histones (Sigma-Aldrich, H9289) were used for targeting chromatin marks and normalization, respectively. Guinea Pig anti-Rabbit IgG (ABIN101961) served as the secondary antibody. Prepared CUT&TAG libraries were sequenced at the Beijing Genomics Institute (BGI) using the DNQSeq 400 platform with a 100+10 base pair paired-end sequencing strategy.

The CUT&TAG sequencing data underwent quality assessment using FastQC-0.12.1. Subsequently, FASTQ files were trimmed using TrimGalore-0.6.10 with options ’--stringency 3’, ’- q 20’, ’--paired’, and ’--length 32’. Alignment of the trimmed reads was performed against the TAIR10 reference genome using Bowtie2-2.5.2 with conditions ’-I 10’, ’-X 700’, ’--no-unal’, ’--no-mixed’, ’--no-discordant’, ’--phred33’, ’--local’, and ’--very-sensitive-local’. The resulting SAM files were converted to sorted BAM files and indexed using Samtools-1.19.2. The bamCoverage function from Deeptools (Ramírez et al., 2016) was used to generate IGV bigwig files, and bigwigCompare was used to generate normalized H3K27me3/H3 IGV bigwig files. MACS2 was used for peak calling and peak annotation.

ComputeMatrix and plotHeatmap functions from deepTools were employed to create heatmaps. K-means clustering was performed using the ’--kmeans’ condition in the plotHeatmap function. GO enrichment analysis was carried out using clusterProfiler (Wu et al., 2021). Network analysis was done using STRING (Szklarczyk et al., 2019). The datasets utilized in this study have been deposited in the NCBI database under reference PRJNA1106003.

## Acknowledgments

We thank Kerstin Zander for technical assistance. We thank Jenny Russinova for providing several reporter lines for BR biosynthesis genes as well as BR mutants, Kerstin Kaufmann for JMJ mutants and REF6 reporter, Juthamas Chaiwanon for *det2*, Sunghwa Choe for *dwf4-5D* and Jie Song for the JMJ13 reporter and 35S::ELF6 line.

This work was funded by the Max Planck Society and by grant number 421178202 of the German Research Foundation (DFG) to DDF, and by the Human Frontier Science Program (LT000162/2018-L) to HS.

## Author contributions

RP, RBL, GYL and DDF designed the study. RP, RBL, GYL, SE, PF and HS performed the experiments. GDT-DL and HB established the CUT&TAG protocols and analysis pipelines. RP, RBL, GYO and SE analyzed the data. RP and DDF wrote the first draft of the manuscript, and all authors contributed to and approved the final version.

## Statements

The authors declare that they have no conflict of interest.

