## Supplement for "BRI1-mediated removal of seed coat H3K27me3 marks is a brassinosteroid-independent process"

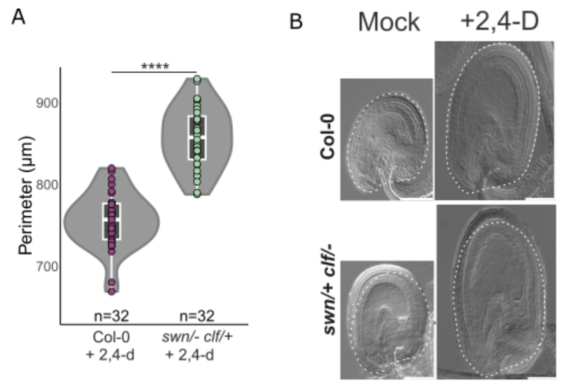


C

| Name | Locus | Known Histone Targets | Expression in the Seed Coat |
|---|---|---|---|
| JMJ11/ELF6 | AT5G04240 | H3K27me2/3 | |
| JMJ12/REF6 | AT3G48430 | H3K27me2/3; H3K36me2/3; H3K4me2/3 | Not Determined |
| JMJ13 | AT5G46910 | H3K27me2/3 | |
| JMJ14 | AT4G20400 | H3K4me1/2/3 | |
| JMJ15/MEE27 | AT2G34880 | H3K4me3 | |
| JMJ16 | AT1G08620 | H3K4me1/2/3 | |
| JMJ17 | AT1G63490 | H3K4me1/2/3 | |
| JMJ18 | AT1G30810 | H3K4me2/3 | |
| JMJ19 | AT2G38950 | H3K4me3 | |
| JMJ20 | AT5G63080 | H4R3me2 | |
| JMJ21 | AT1G78280 | Not Determined | |
| JMJ22 | AT5G06550 | H4R3me2 | |
| JMJ24 | AT1G09060 | H3K9me2 | |
| JMJ25/IBM1 | AT3G07610 | H3K9me1/2 | |
| JMJ26 | AT1G11950 | Not Determined | |
| JMJ27 | AT4G00990 | H3K9me1/2 | |
| JMJ28 | AT4G21430 | H3K9me1/2 | |
| JMJ29 | AT1G62310 | H3K9me1/2; H3K4me3 | |
| JMJ30 | AT3G20810 | H3K27me2/3; H3K36me2/3 | |
| JMJ31 | AT5G19840 | Not Determined | |
| JMJ32 | AT3G45880 | H3K27me2/3 | |

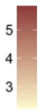


**Figure S1.** (**A**) Seed size differences of auxin-induced autonomous seeds in WT and *swn clf/+* at 3 days after auxin treatments (3 DAT). Representative photos are seen in (**B**). Scale bars indicate 50 µm. (**C**) List of known JMJ-type demethylases with their putative substrates and expression levels in seed coats of seeds at the pre-globular embryo stage. The bar on the right-hand side represents the relative expression values, as previously determined (Belmonte et al., 2013).

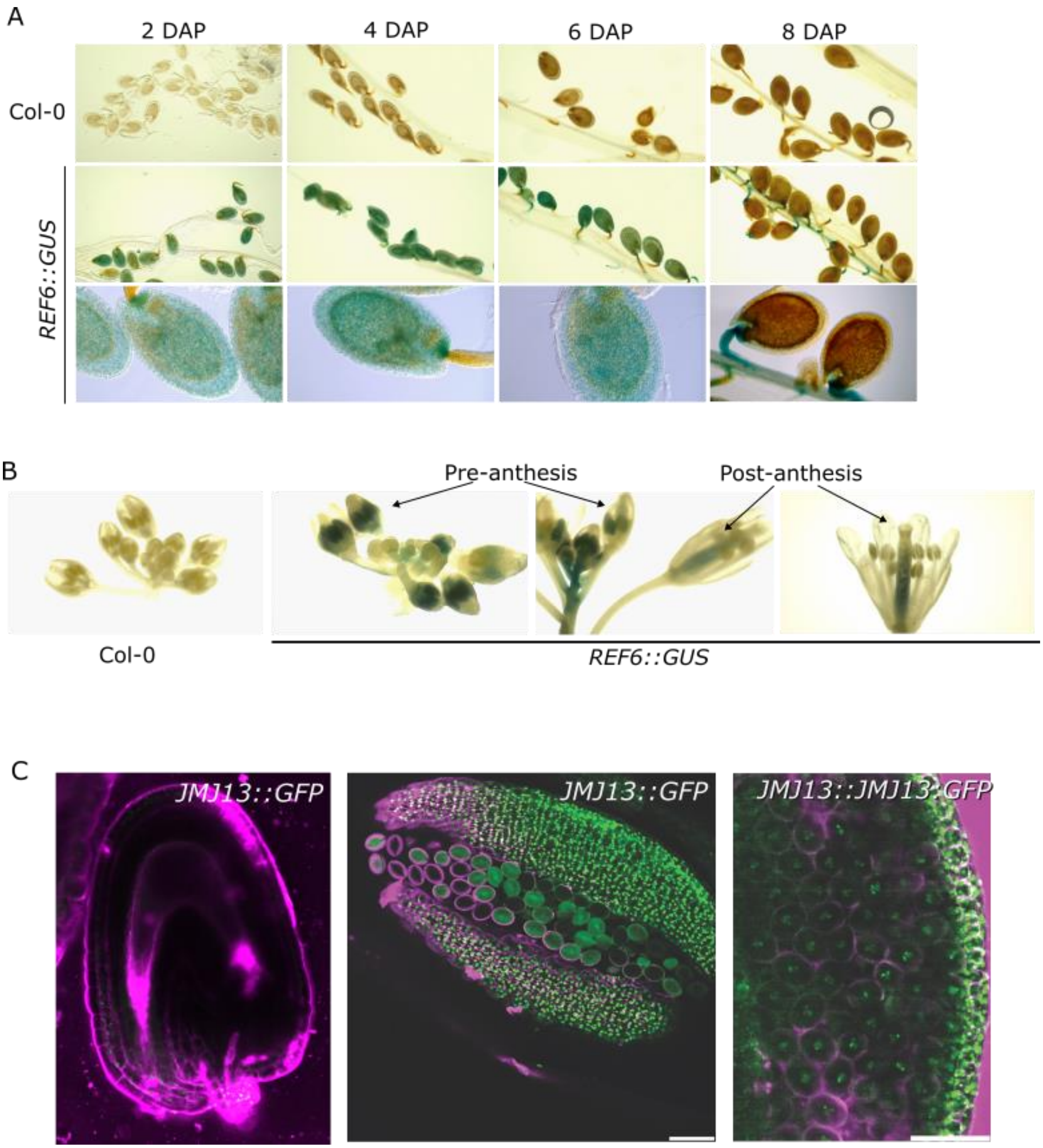


**Figure S2. Tissue specific expression of JMJ demethylases.** (**A-B**) Expression of REF6 as determined by beta-glucuronidase activity in developing seeds (**A**) and flowers (**B**). Untransformed plants (Col-0) were used as negative control. (**C**) Expression of *JMJ13* in seeds and anthers as determined by *JMJ13::GFP and JMJ13:JMJ13:GFP*. Scale bars indicate 50 µm.

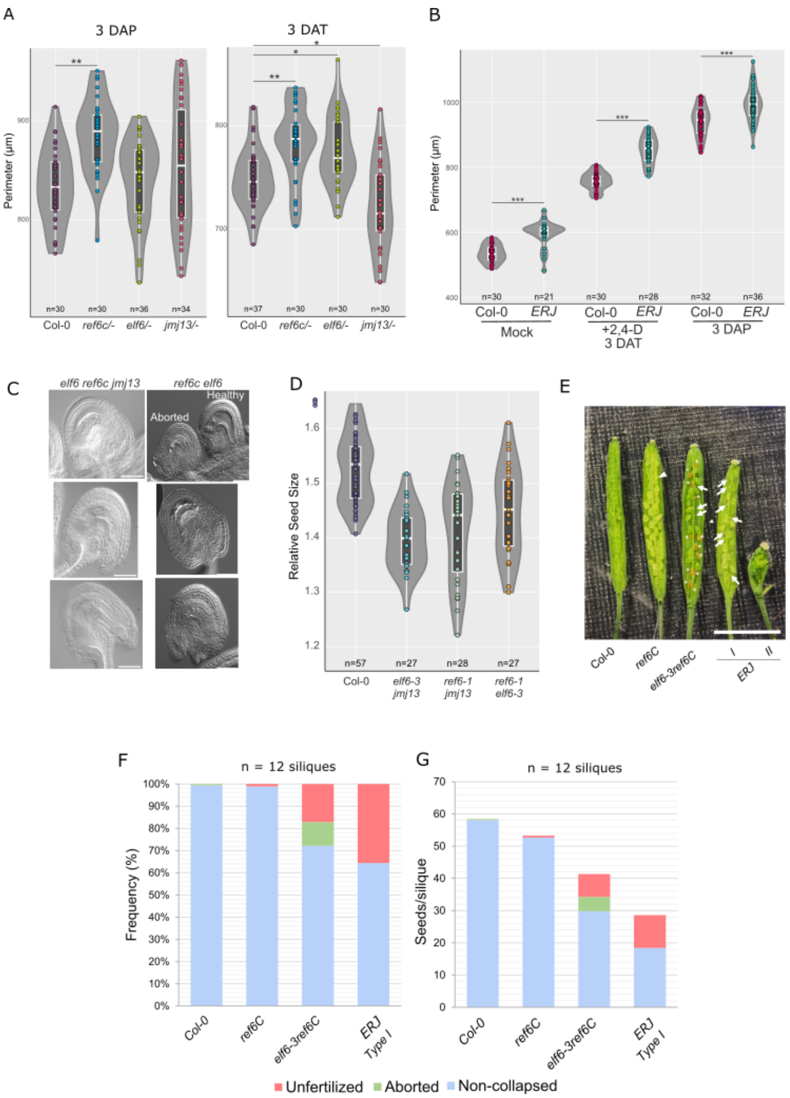


**Fig. S3.** (**A**) Size of fertilized (3 DAP) and auxin-induced seeds (3 DAT) for the single *jmj* mutants and respective WT. (**B**) Size of unfertilized ovules (Mock), autonomous seeds (3 DAT) and sexual seeds (3 DAP) of the *ERJ* mutant and respective WT. Differences are significant for p<0.01 (*), p<0.001 (**) or p<0.00001 (***) (ANOVA). (**C**) Aborted ovules of *ERJ* (left) and *ref6c elf6* (right). Scale bars indicate 50 µm. (**D**) Relative size of WT and *jmj* double mutant seeds at 3 DAP. (**E**) Morphology of *ref6c*, *elf6 ref6c and ERJ* siliques. Two types of siliques are obtained in *ERJ:* Type I siliques are more morphologically similar to the WT, while Type II siliques are malformed and often show “carpel-within-carpel” phenotypes (more details in **Fig. S4**). Aborted seeds are indicated by arrows and unfertilized/underdeveloped ovules are indicated with asterisks. Scale bar indicates 5 mm. (**F**) Quantification of seed phenotypes in Col-0, *ref6C*, *elf6-3 ref6C* and *ERJ*, scored as indicated in (**E**). (**G**) Mature seed set of Col-0, *ref6C*, *elf6-3 ref6C* and *ERJ*. Note the extremely reduced seed set of ERJ. For (**F**) and (**G**) only Type I siliques were taken into account. The seeds from twelve independent siliques were counted: three siliques of four different plants each.

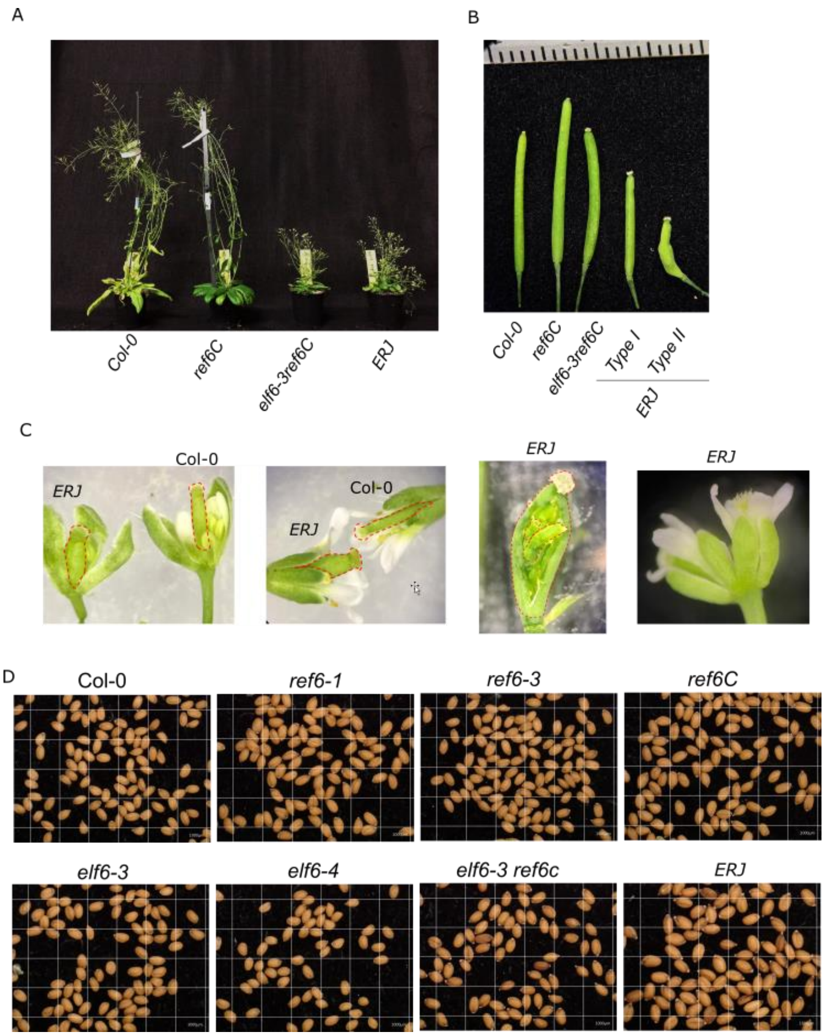


**Fig. S4**. **Vegetative and seed phenotypes of Col-0 and *jmj* mutants.** (**A**) Overall plant morphology of Col-0, *ref6c*, *elf6-3 ref6c* and *ERJ*. Note the dwarfism of *efl6-3 ref6c* and *ERJ*. (**B**) Bent silique phenotype observed in Type II siliques of *ERJ* (also described in **Fig. S3**). (**C**) Floral phenotypes of *ERJ*, showing swollen pistils and pitil-within-pistil phenotypes. None of these phenotypes are observable in the other genotypes, including in *elf6-3 ref6c*. (**D**) Mature seed phenotypes of Col-0, *ref6* and *elf6* single mutants and higher order *elf6-3 ref6c* and *ERJ* mutants.

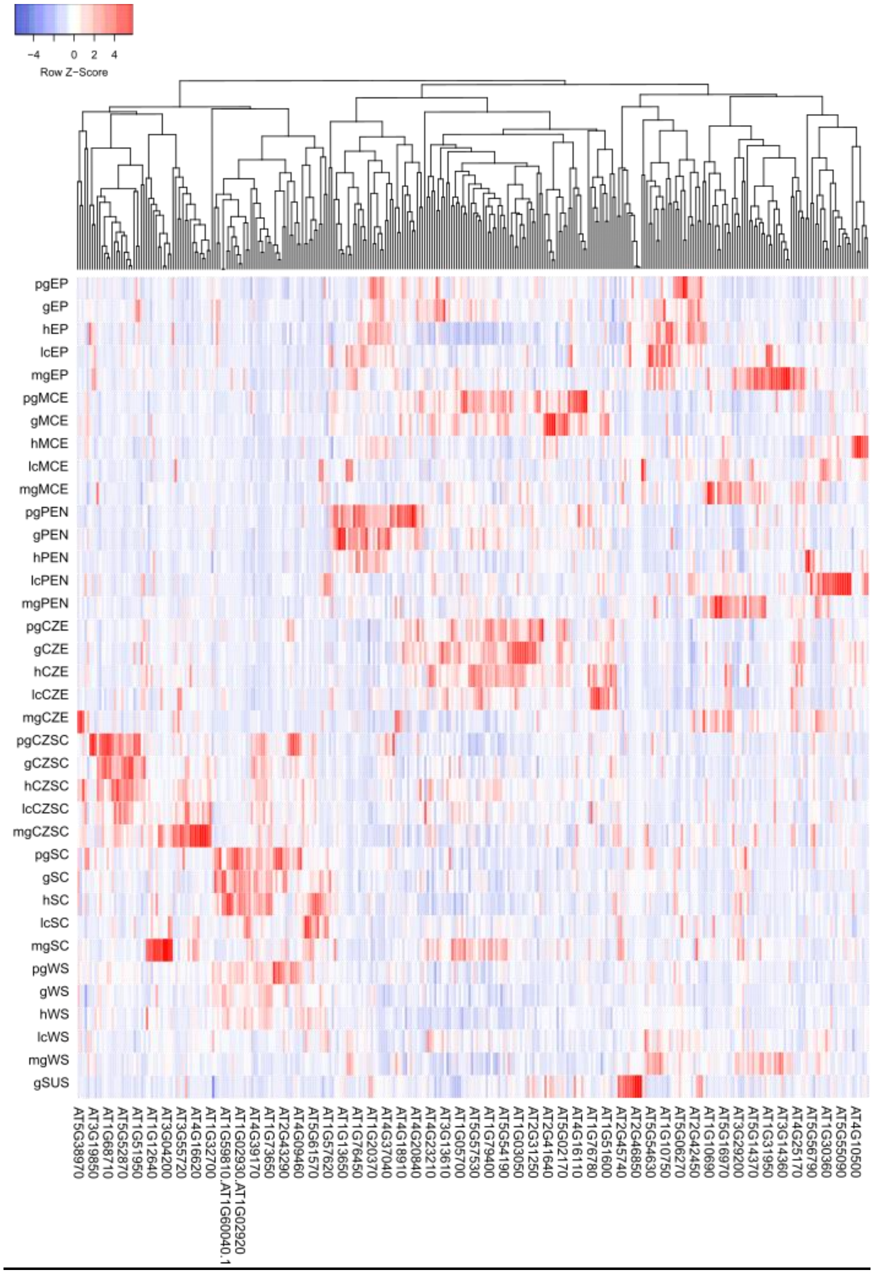


**Fig. S5.** Relative expression of genes carrying REF6 binding sites, specifically expressed in the embryo proper (EP), micropylar endosperm (MCE), peripheral endosperm (PEN), chalazal endosperm (CZE), chalazal seed coat (CZSC), seed coat (SC), whole seed (WS) and suspensor (SUS). Gene expression data is based on (Belmonte et al., 2013). The seed stages indicated are: pg, pre-globular; g, globular; h, heart; lc, linear cotyledon; and mg, mature green. Only genes with at least 4 REF6 binding sites were included in the analysis.

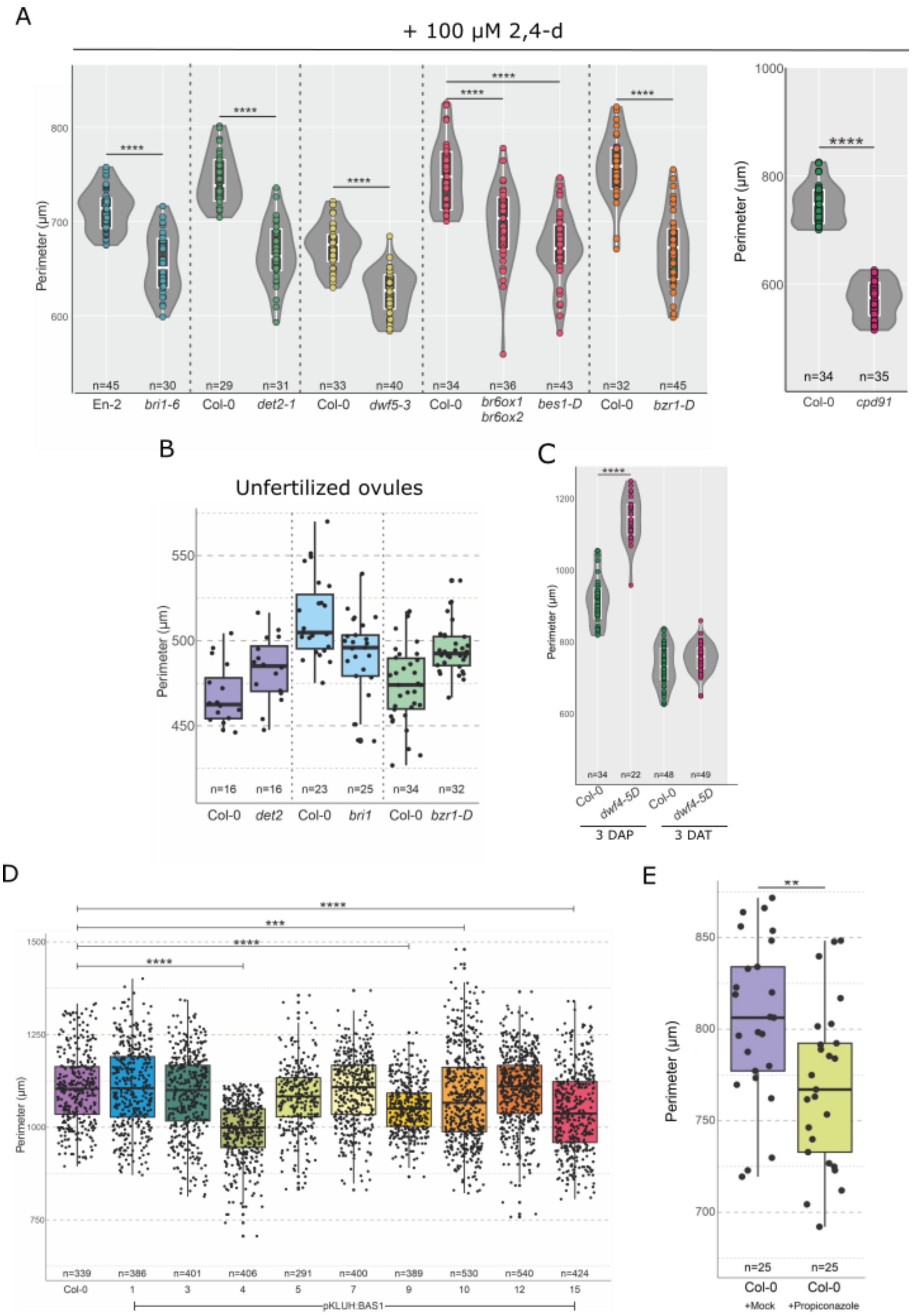


**Fig. S6.** (**A**) Perimeter of auxin-induced autonomous seeds as determined at 3 DAT, for gain- and loss-of-function mutants in BR biosynthesis and signaling. (**B**) Size of unfertilized ovules of Col-0 WT, *det2*, *bri1* and *bzr1-1D*. Differences are not statistically different between the WT and respective mutants. (**C**) (**A**) Perimeter of fertilized and of auxin-induced autonomous seeds as determined at 3 DAP and 3 DAT, respectively, for the BR overproducing mutants *dwf4-5d*. (**D**) Seed perimeter of dry seeds of transgenic lines expressing *KLUH::BAS1*. ** indicates statistical significance for p-value <0.01, and **** for p-value <0.0001 (Anova). (**E**) Seed size at 3 DAP after treatments with the BR inhibitor propiconazole.

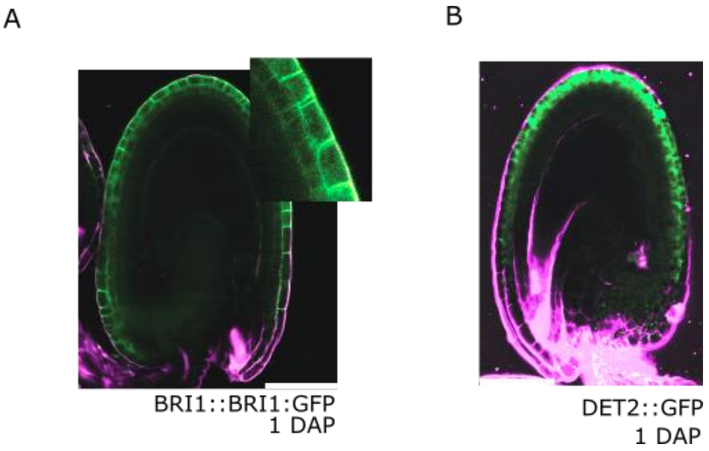


**Fig. S7**. Expression of *BRI1::BRI1:GFP* (**A**) and of *DET2::GFP* (**B**) in seeds at 1 DAP. Magenta is propidium iodide (PI). Scale bars indicate 50 µm.

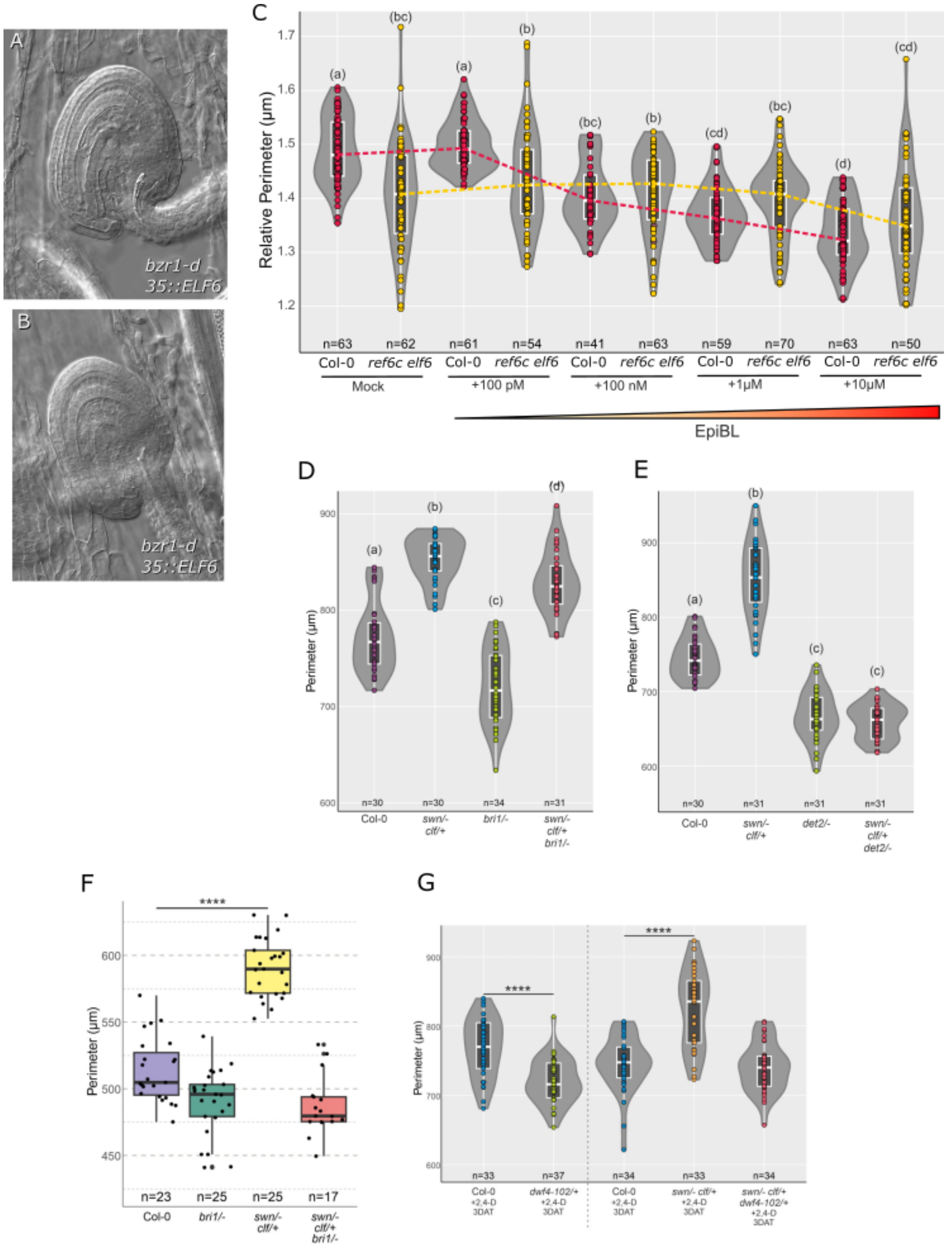


**Fig. S8**. (**A-B**) Examples of malformed ovules as obtained in lines expressing *CaMV35S::ELF6 bzr1-D*. No such phenotypes were observed in the single mutants. Scale bars indicate 50 µm. (**C**) Perimeter of auxin+epi-BL treated WT (Col-0) and *ref6 elf6* seeds at 3 DAT. The letters indicate statistical significance for p-value <0.05 (Anova). (**D**) Perimeter of autonomous seeds after auxin application in WT, *swn clf*/+ and either *bri1-6* (**C**) or *det2-1* (**D**), and respective higher order mutants. The letters indicate statistical significance for p-value <0.05 (Anova). (**E**) Ovule size of WT, *bri1*, *swn clf*/+ and respective double mutant at 2 DAE. **** indicates p-value <0.0001 (Pairwise t-test). (**F**) Perimeter of autonomous seeds after auxin application in WT, *swn clf*/+, *dwf4-102* and respective triple mutant. **** indicates p-value <0.0001 (Anova).

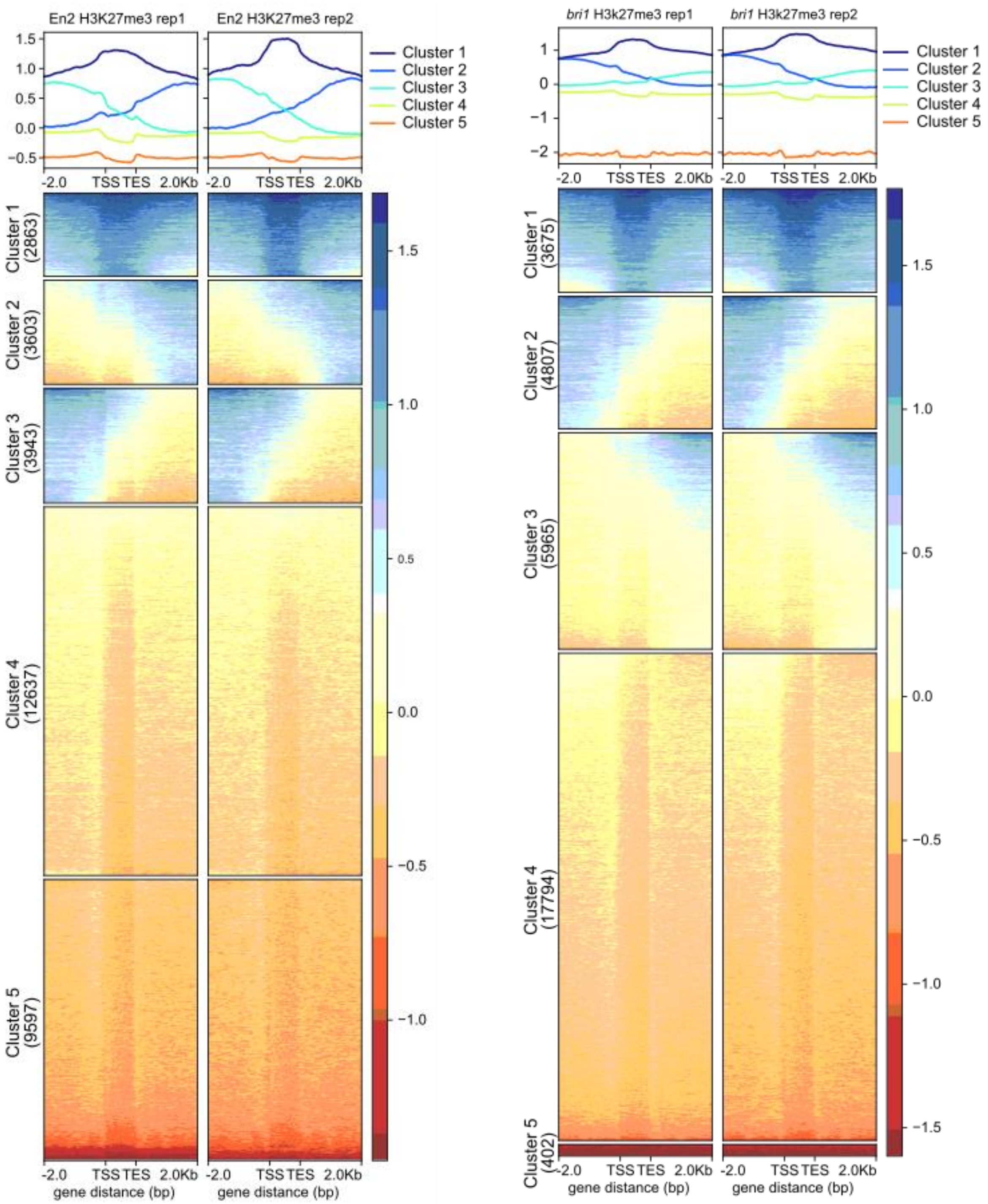


**Fig. S9.** Clustering analysis of H3K27me3 profiles in WT En-2 and in *bri1*. The two replicates are shown for each genotype. The metagene plots on top indicate the average enrichment of H3K27me3 over gene regions for each of the Clusters indicated below. The colors indicate relative enrichment. TSS - transcription start site, TES - transcription end site.

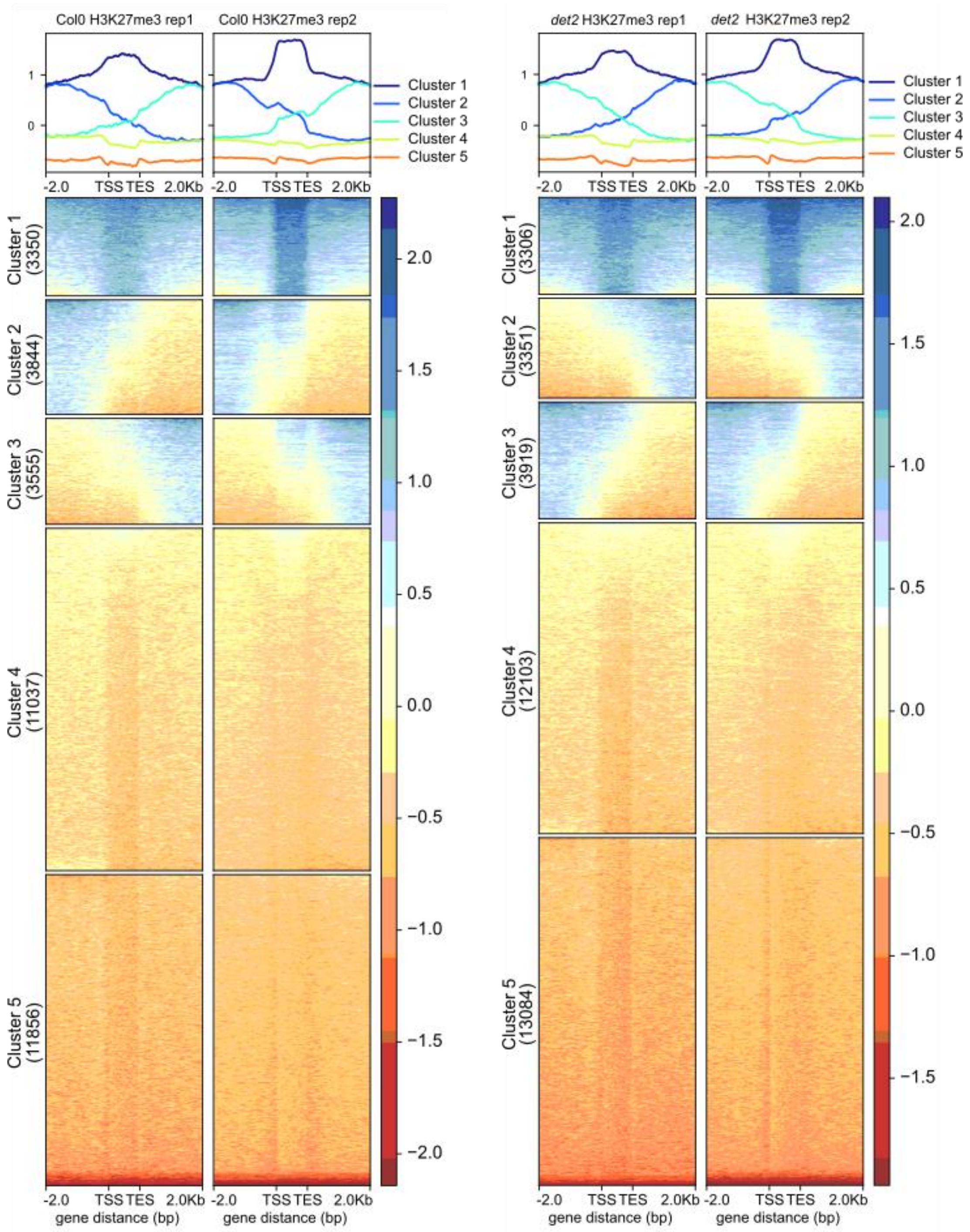


**Fig. S10.** Clustering analysis of H3K27me3 profiles in WT Col-0 and in *det2*. The two replicates are shown for each genotype. The metagene plots on top indicate the average enrichment of H3K27me3 over gene regions for each of the Clusters indicated below. The colors indicate relative enrichment. TSS - transcription start site, TES - transcription end site.

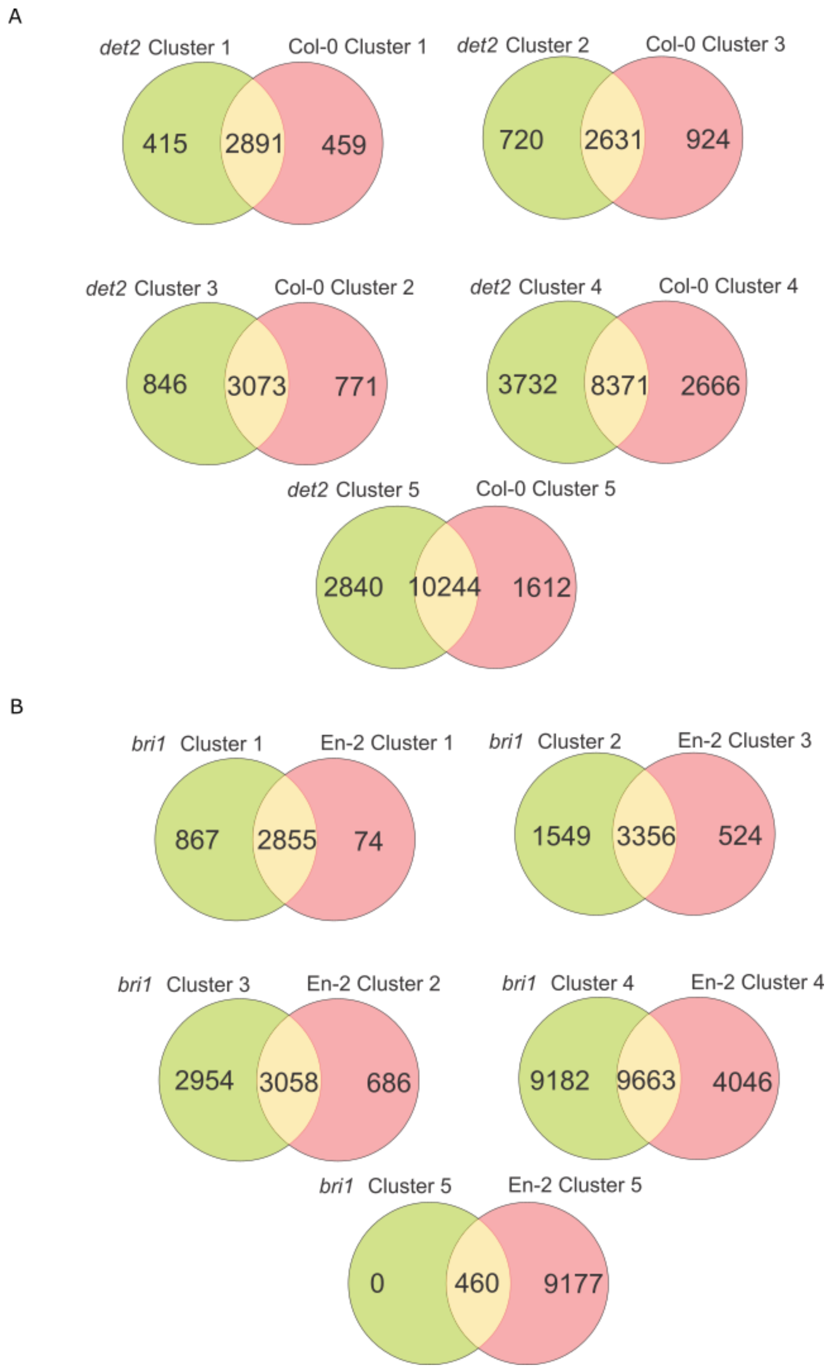


**Fig. S11.** Overlaps between the number of genes labeled with H3K27me3 between *det2* and Col-0 (**A**) and *bri1* and En-2 (**B**) Clusters. Similar clusters were compared between the pairs of genotypes. Note that WT Clusters 2 were compared to the mutant Clusters 3. This is because WT Clusters 2 represent genes labeled in the promoter region, while genes labeled in the terminator are in Cluster 3. In both mutants this is the opposite.Therefore we decided to compare the most similar clusters between the genotypes.

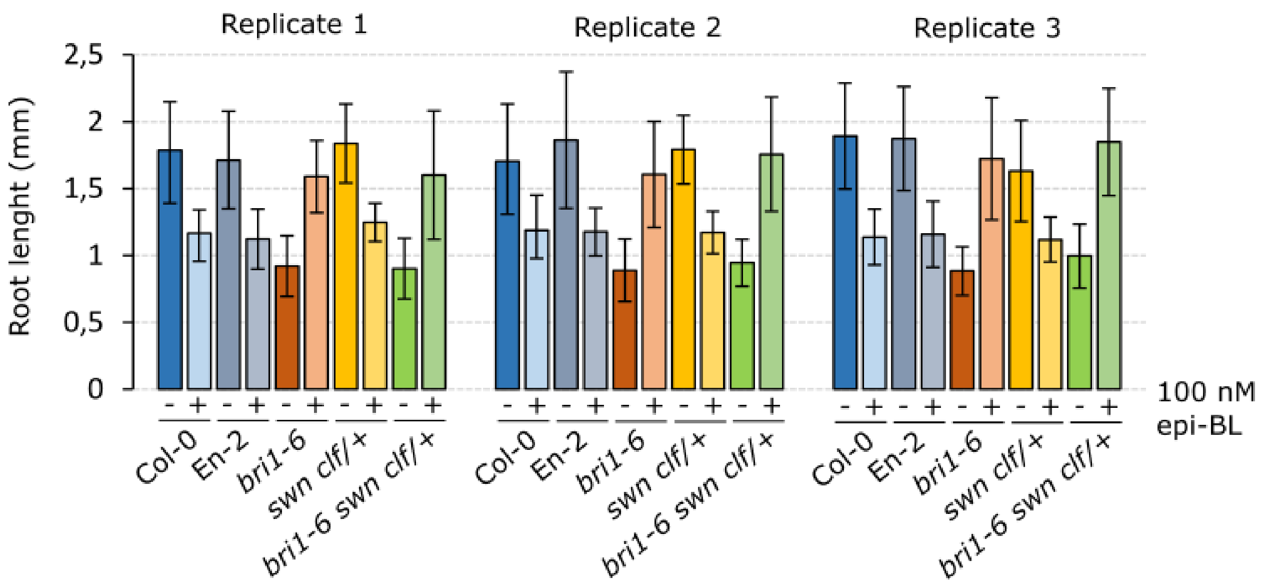


**Fig. S12.** BR root sensitivity assay. Three independent replicates are shown, grown either in ½ MS plates or ½ MS supplemented with 100 nM epi-BL. The roots were measured at 7 days after germination. The bars indicate standard deviation.

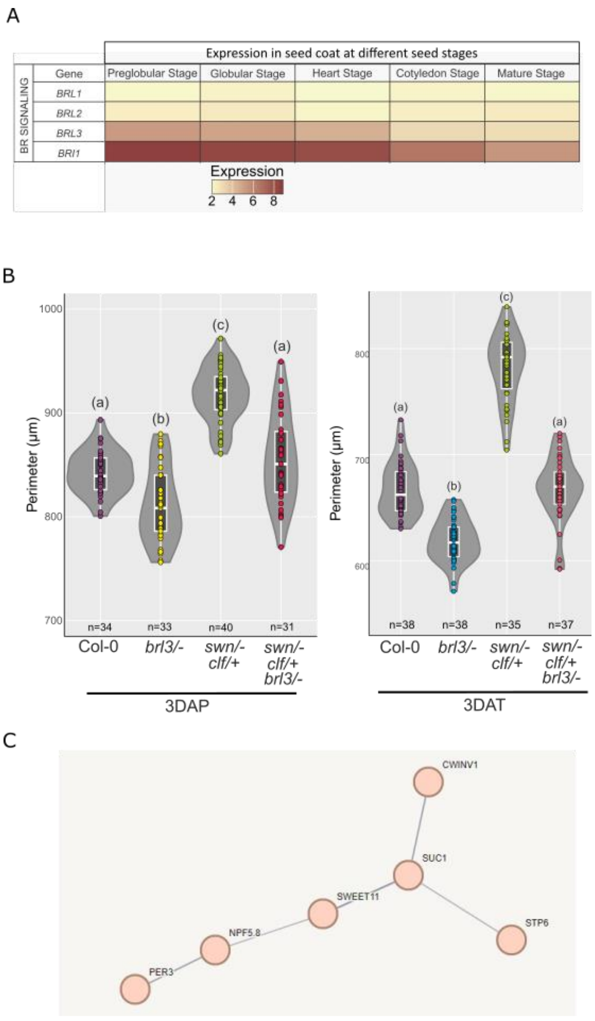


**Fig. S13.** (**A**) Expression of *BRL* genes in different stages of seed development. (**B**) Seed perimeter of Col-0, *brl3*, *swn clf*/+, and respective triple mutant, at 3 DAP and 3 DAT. The letters indicate statistical significance for p-value <0.05 (Anova). (**C**) Cluster of sugar transport related genes differentially methylated by H3K27me3 in *brl3* mutants.

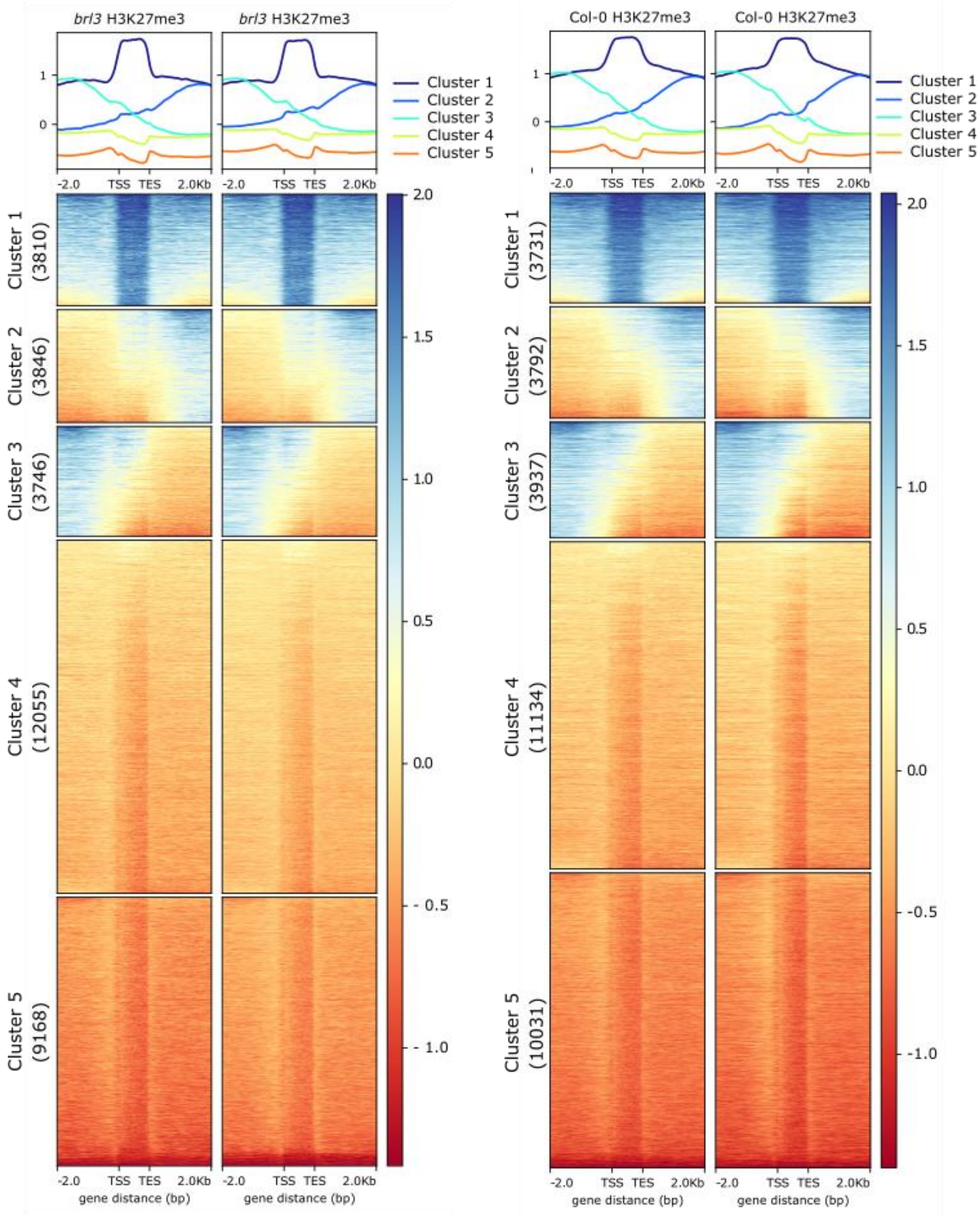


**Fig. S14.** Clustering analysis of H3K27me3 profiles in WT Col-0 and in *brl3*. The two replicates are shown for each genotype. The metagene plots on top indicate the average enrichment of H3K27me3 over gene regions for each of the Clusters indicated below. The colors indicate relative enrichment. TSS - transcription start site, TES - transcription end site.

**Table S1**: Primers used for genotyping and cloning.

| Primer | Genotype | Sequence | Information |
|---|---|---|---|
| p2_1 | *jmj13* | TCAAGTTCCATTTGCTGCTTC | WT p2_1 & p2_2 |
| p2_2 | *jmj13* | GGAATGTTTGCTGACACTTGG | Mut p2_2 &p2_3 |
| p2_3 | *jmj13 tdna (GABIKAT)* | TGGTTCACGTAGTGGGCCATCG | |
| p2_4 | *elf6-3* | ACGTCAATGCGGTAATCATTC | WT p2_4 & p2_5 |
| p2_5 | *elf6-3* | TTTGCAGATCCCATTGCTTAC | Mut p2_5 & p2_8 |
| p2_6 | *ref6-1* | TCATATACAAGGCGTTCGGTC | WT p2_6 & p2_7 |
| p2_7 | *ref6-1* | CAGTTGCAACTCTGGAGAAGG | Mut p2_7 & p2_8 |
| p2_8 | *SALK T-DNA LBb1.3* | ATTTTGCCGATTTCGGAAC | genotyping SALK lines |
| p2_9 | *ref6c* | GGTTGCTCCAGAGTTCAGACC | Mutant band smaller than WT |
| p2_10 | *ref6c* | CATGGTCCTCCACATGCCAAGC | Mutant band smaller than WT |
| p2_11 | *JMJ13 promoter gateway FP* | GGGGACAAGTTTGTACAAAAAAGCAGGCTTCttaaaaagggaataggagatttaag | cloning JMJ13 transcriptional reporter |
| p2_12 | *JMJ13 promoter gateway RP* | GGGGACCACTTTGTACAAGAAAGCTGGGTTATCATTCCGTAATCTATTATCTCTG | cloning JMJ13 transcriptional reporter |
| p2_13 | *JMJ13 promoter sequencing* | cccaagattttaagcaaagta | sequencing JMJ13 transcriptional reporter |
| p3_15 | *dwf4-102* | GGCAGCTCCTACGTCATTAAAG | WT p3_15+p3_16 |
| p3_16 | *dwf4-102* | CCGGACATGAGACTTCTTCTG | Mut p3_16+p3_1 |
| p3_17 | *dwf4-5D* T-DNA | aaaggctatcgttcaagatgcc | |
| p3_18 | *dwf4-5D* | AGCGTGTAACCATCTGCAAC | WT p3_18+p3_19 |
| p3_19 | *dwf4-5D* | GTCAAACCCGTAGACTTTTGGC | WT p3_17+p3_19 |
| p3_20 | *bzr-1D* | CTAACTGGGAATCTATCGC | To be cut by HpaII |
| p3_21 | *bzr-1D* | TCAACCACGAGCCTTCC | WT (127bp + 310 bp) Mu (437 bp) |
| p3_22 | *brl3* | CCAGTGAACTCGTTTGAGCTC | WT p3_22+p3_23 |
| p3_23 | *brl3* | TTTATCGAACACTTTGTGGGC | Mut p3_23+p3_1 |
| p3_24 | *swn-3* | CGTTTCCGAGGATGTCATTGTG | WT p3_24+p3_25 |
| p3_25 | *swn-3* | TGGAACTTTTGAGTGGCTAGAGGTG | WT p3_25+p3_26 |
| p3_26 | SALK T-DNA LBb1.1 | GCGTGGACCGCTTGCTGCAACT | genotyping SALK lines |
| p3_27 | *clf9* | tgggttcgtttaggaaccatt | WT p3_27+p3_28 |

| | | | |
|---|---|---|---|
| p3_28 | *clf9* | ccagcataacagttgacatagca | WT p3_26+p3_28 |
| p3_29 | DET2 CDS | GGGGACAAGTTTGTACAAAAAAGCAGGCTatggaagaaatcgccgataaaac | cloning DET2 in rescue lines |
| p3_30 | DET2 CDS | GGGGACCACTTTGTACAAGAAAGCTGGGTtcagtacacaaaaggaataacagct | cloning DET2 in rescue lines |
| p3_31 | DET2 seq | TGAGCGAAACCCTATAAGAACCCT | Genotyping DET2 rescue lines p3_29+p3_31 |
| p3_32 | BRI1 CDS | GGGGACAAGTTTGTACAAAAAAGCAGGCTATGAAGACTTTTTCAAGCTTCTTTCT | cloning BRI1 in rescue lines |
| p3_33 | BRI1 CDS | GGGGACCACTTTGTACAAGAAAGCTGGGTTCATAATTTTCCTTCAGGAACTTCT | cloning BRI1 in rescue lines |
| p3_34 | BRI1 seq | GTGGGTTTGGAGATGTTTAC | Sequencing BRI1 in rescue lines |
| p3_35 | BRI1 seq | GGTAAACGGCCAACGGATT | Genotyping BRI1 rescue lines p3_33+p3_35 |
| p3_36 | GUSplus CDS | gcgggatatcactagATGGTAGATCTGAGGGTAAATTTCT | Cloning GUSplus in pB7WG |
| p3_37 | GUSplus CDS | ttgaacgatcctgcaTCAGTTCTTGTAGCCGAAATCTGGA | Cloning GUSplus in pB7WG |
| p3_38 | REF6 promoter | ataattcgagggtacATGAAGAGTTGAGTGATGACACG | Cloning REF6 promoter in reporter lines |
| p3_39 | REF6 promoter | cgtcgggccctctagATCTCTCTCTCTCTCTCACACACACAGGG | Cloning REF6 promoter in reporter lines |
| p3_40 | ELF6 promoter | ataattcgagggtacATTGCTCGTTTAACAAGACCGTG | Cloning ELF6 promoter in reporter lines |
| p3_41 | ELF6 promoter | cgtcgggccctctagCTTAAATTCCCAATTCCGTAAGACC | Cloning ELF6 promoter in reporter lines |
| p3_42 | BRL3 promoter | GAGCTCGTTCGGCGCAAATACT CDAC | Cloning BRL3 reporter construct |
| p3_43 | BRL3 promoter | GCTAGCGTTATTAGCCCACAAAGTGTTCGA | Cloning BRL3 reporter construct |
